# Alternatively Spliced Landscape of PPARγ mRNA in Podocytes is Distinct from Adipose Tissue

**DOI:** 10.1101/2022.09.12.507605

**Authors:** Claire Bryant, Amy Webb, Alexander S Banks, Dawn Chandler, Rajgopal Govindarajan, Shipra Agrawal

## Abstract

Podocytes are highly differentiated epithelial cells, and their structural and functional integrity is compromised in a majority of glomerular and renal diseases, leading to proteinuria, chronic kidney disease, and kidney failure. Traditional agonists (e.g., pioglitazone) and selective modulators (e.g., GQ-16) of peroxisome-proliferator-activated-receptor-γ (PPARγ) reduce proteinuria in animal models of glomerular disease and protect podocytes from injury via PPARγ activation. This indicates a pivotal role for PPARγ in maintaining glomerular function through preservation of podocytes distinct from its well-understood role in driving insulin sensitivity and adipogenesis. While its transcriptional role in activating adipokines and adipogenic genes is well-established in adipose tissue, liver and muscle, understanding of podocyte PPARγ signaling remains limited. We performed a comprehensive analysis of PPARγ mRNA variants due to alternative splicing, in human podocytes and compared with adipose tissue. We found that podocytes express the ubiquitous PPARγ Var 1 (encoding γ1) and not Var2 (encoding γ2), which is mostly restricted to adipose tissue and liver. Additionally, we detected expression at very low level of Var4, and barely detectable levels of other variants, Var3, Var11, VartORF4 and Var9, in podocytes. Furthermore, a distinct podocyte vs adipocyte PPAR-promoter-response-element containing gene expression, enrichment and pathway signature was observed, suggesting differential regulation by podocyte specific PPARγ1 variant, distinct from the adipocyte-specific γ2 variant. In summary, podocytes and glomeruli express several PPARγ variants, including Var1 (γ1) and excluding adipocyte-specific Var2 (γ2), which may have implications in podocyte specific signaling and pathophysiology. This suggests that new selective PPARγ modulators can be potentially developed that will be able to distinguish between the two forms, γ1 and γ2, thus forming a basis of novel targeted therapeutic avenues.

## 1. Introduction

Thiazolidinediones (TZDs) or peroxisome proliferator activated receptor γ (PPARγ) agonists such as pioglitazone, directly protect podocytes from injury as demonstrated in podocyte culture *in vitro* studies [1–5], and reduce proteinuria and glomerular injury in various animal models of glomerular disease, as reported in preclinical *in vivo* studies [5–13]. Moreover, these beneficial effects of PPARγ have been shown to be mediated by activation of podocyte PPARγ as demonstrated by elegant studies using podocyte specific *Pparg* knock out (KO) mouse model [7, 10]. This indicates a pivotal role for PPARγ in maintaining glomerular function through preservation of podocytes [1, 5–7, 10]. Podocytes are highly differentiated epithelial cells in the kidneys, whose structural and functional integrity is critical for the maintenance of glomerular filtration barrier [14–16]. Accordingly, their dysfunction or loss is the initiating and progressing characteristic factor in a vast majority of renal diseases, leading to chronic kidney disease and kidney failure. Glomerular diseases characterized by high proteinuria manifest as nephrotic syndrome (NS) which is often associated with co-morbidities such as hypoalbuminemia, hypercholesterolemia, edema, and hyper-coagulopathy [17–22]. Recently, our group has further demonstrated that selective modulation of PPARγ with a partial agonist, GQ-16, provides high efficacy in reducing proteinuria, as well as NS-associated comorbidities in an experimental model of nephrotic syndrome [23]. Thus, a multitude of evidence shows that targeting the podocyte PPARγ pathway offers an attractive therapeutic strategy in NS. While PPARγ has an established role in driving many adipogenic and lipid metabolizing genes mainly in adipose tissue, liver, and muscle, the understanding of molecular pathways regulated by PPARγ in podocytes and glomeruli remains limited [5–7, 9, 10, 24–28]. Furthermore, it is well-known that PPARγ exists in two major isoforms, γ1 and γ2, which are a result of different promoter usage as well as alternative splicing (AS) [29–31]. As a result, γ2 contains 30 additional amino acids at its N terminal, despite a longer mRNA sequence towards the 5’UTR of γ1. While γ1 is more widely expressed, γ2 is mostly restricted to adipose tissue and liver. Few other forms have been discovered, including some with dominant negative function [32–37]. Moreover, cell-specific expression of PPARγ variants has been demonstrated to play a differential role in downstream gene expression pattern in adipocytes (expressing the γ2 variant) vs the macrophages (expressing the γ1 variant) [38]. However, the specific variant or isoform(s) expression of *PPARγ* and the role of its AS in podocytes and glomerular disease is unexplored. In this study, we sought out to answer this critical gap in literature by hypothesizing that alternative splice variants of PPARγ expressed in podocytes are distinct from adipocytes. To address this hypothesis, we performed a comprehensive analysis of different AS variants of PPARγ in human podocytes and studied the expression of γ1 vs γ2 variant in podocyte vs adipose tissue in detail.

## 2. Materials and Methods

### Podocyte Cell Culture and Treatments

Immortalized human podocytes were cultured in RMPI 1640 (Corning, Tewksbury, MA) supplemented with 10% fetal bovine serum (FBS) (Corning, Tewksbury, MA), 1% 100X Penicillin Streptomycin L-Glutamine (PSG) (Corning, Tewksbury, MA), and 1% Insulin-Transferrin-Selenium-Ethanolamine (ITS-X) (Gibco, Gaithersburg, MD) [39]. Proliferating podocytes were cultured in a humidified atmosphere with 5% CO2 at 33ºC, and the media was changed twice per week and cells were passaged at ~70%-80% confluence. Differentiation was induced by placing the cells at 37ºC under the same atmospheric conditions, for 14 days. Differentiated cells were treated with puromycin aminonucleoside (PAN) (Sigma-Aldrich, St. Louis, MO) at 25 ug/ml or with pioglitazone (Alfa Aesar, Tewksbury, MA) at 10 μM. Control cells received treatment of vehicle dimethyl sulfoxide (DMSO) (Sigma-Aldrich, St. Louis, MO). Cells were harvested after 24 hours and total RNA isolated.

### Kidney, Liver and Adipose Tissues

This study was approved by the Institution Animal Care and Use Committee at Nationwide Children’s Hospital and the experiments were performed according to their guidelines. Male Wistar rats weighing ~150-200 g (Envigo, Indianapolis, IN) were purchased, acclimated for 3 days, and housed under standard conditions. They were sacrificed to collect kidneys, liver, and white adipose tissue (WAT) epididymal fat. Tissues were flash frozen in liquid nitrogen prior to RNA isolation. Glomeruli were isolated from harvested kidneys using the sequential sieving method as previously described [23]and total RNA was isolated. Commercially available human and rat kidneys total RNA was purchased from Zyagen (San Diego, CA).

### RNA Isolation

Rat epididymal fat and liver tissue samples were lysed in RNA extraction buffer (RLT) from the RNeasy kit (Qiagen, Germantown, MD) with stainless-steel disruption beads for 4 minutes at 30.0 Hz using the Qiagen TissueLyser (Germantown, MD). Total RNA was immediately isolated from the lysate using the RNeasy kit (Qiagen, Germantown, MD), following the manufacturer’s instructions. Total RNA was extracted from rat glomeruli tissue and human podocyte samples using the mirVanaTM Isolation Kit (Invitrogen, Carlsbad, CA), according to the manufacturer’s instructions. RNA yield and purity was checked prior to downstream applications by measuring the absorbance at 260, 280, and 230 nm with a nanodrop spectrophotometer (ThermoFisher, Waltham, MA) and by calculating appropriate ratios (260/280, 260/230).

### DNase Treatment, cDNA Synthesis, and PCR

Total RNA was DNase treated according to manufacturer’s instructions (Zymo Research, Irvine, CA). 500 ng - 1μg DNase-treated RNA was reverse transcribed using the iScript cDNA Synthesis Kit (Bio-Rad, Hercules, CA), according to the manufacturer’s instruction and the resulting cDNA was used for reverse transcription-polymerase chain reaction (RT-PCR) using HotStarTaq Plus Master Mix Kit (Qiagen, Germantown, MD). The PCR conditions used were as follows: 95°C for 5 minutes, 40 cycles [95°C for 30 seconds, 50-60°C (depending on the primer pair) for 30 seconds, 72°C for 30 seconds], hold at 4°C. Housekeeping gene *RPL6* was amplified only for 28 cycles to prevent over-saturation. The PCR-products were separated on a 1.5-2.5% agarose gel along with molecular weight ladder (ThermoFisher, Waltham, MA), stained with 0.5µg/mL ethidium bromide (Research Products International, Mount Prospect, IL), and images were captured using the Chemidoc (Bio-Rad, Hercules, CA) equipment. Densitometry was performed using ImageJ Software (National Institutes of Health, Bethesda, MD) and band density was calculated by subtracting background and normalizing to *RPL6*. Quantitative RT-PCR (qRT-PCR) was performed on samples using SYBR green (Bio-Rad, Hercules, CA) on the Applied Biosystems 7500 Real-Time PCR System (Waltham, MA). The PCR conditions were 95°C for 10 minutes, 40 X (95°C for 15 seconds, 50-60°C for 1 minute) followed by a melt curve to ensure specific products. Analysis was performed using the ∆∆Ct method [40] with normalization to *RPL6*.

### Primer Design and Synthesis

Primers were custom designed to detect various PPARγ variants for alternatively spliced mRNA products using the Reference Sequence Database (National Center for Biotechnology Information, Bethesda, MD) and confirmed for the potential binding transcripts in the annotated database using the Primer Blast program (National Center for Biotechnology Information, Bethesda, MD) (Table 1, Figure 1A). For variants with exon skipping, primers were designed to span the exon-exon junction to create a specific primer pair that would exclusively amplify the desired mRNA variant.

**Figure 1.**
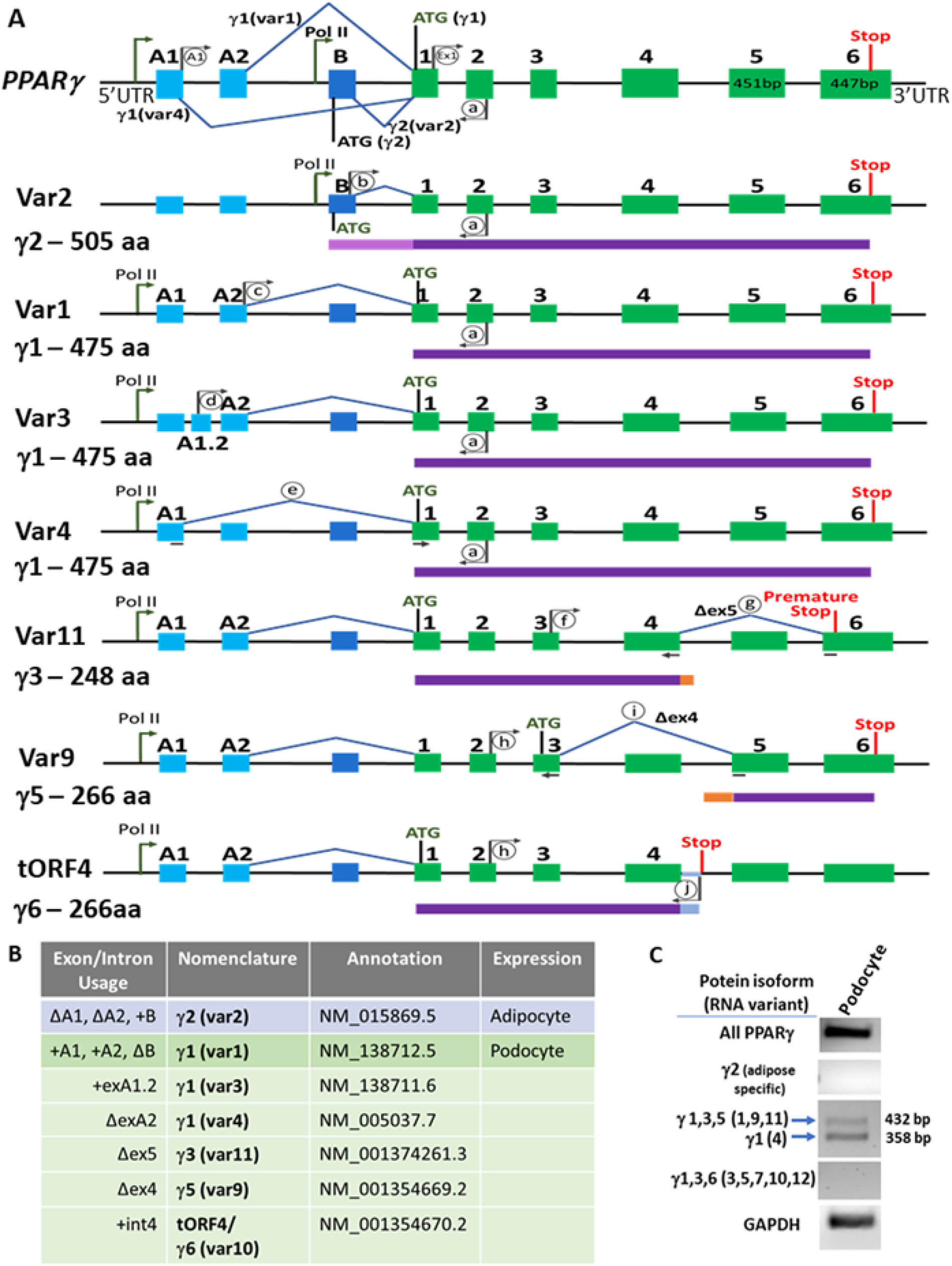
Alternatively Spliced Variants and Isoforms of PPARγ in Podocytes. (A) The schematic depicts the exon and intron usage and the transcription and translation start and stop sites for various PPARγ variants. The major isoforms of PPARγ are isoforms γ1 and γ2, which encode for proteins containing 475 and 505 amino acids, respectively. The most common human annotated variants 1 (var1: NM_138712) and 2 (var2: NM_015869) encode isoforms γ1 and γ2 (depicted in the top-most PPARγ illustration). These two variants are transcribed by different promoter usage by RNA Polymerase II (depicted in tall arrows) and alternative splicing (depicted by exon skipping). PPARγ1 variant 1 includes exons A1 and A2, a start codon (ATG) on exon 1, and γ2 variant 2 excludes exons A1 and A2, but includes exon B and a start codon in the same exon B. Thus, γ1 protein product is 30 amino acids shorter than γ2. The less known and understood variants are represented below, which include Variants 3, 4, 11, 9 and tORF4/Var 10. These are depicted with A1 and A2 usage in the schematic to emphasize those likely to be expressed in podocytes, although their adipocyte counterparts with exon B usage at the 5’ UTR also exist. A1 and A2 exons are shown in light blue, exon B in dark blue, internal exons are in green boxes, introns are black lines, spliced out exons are marked with blue lines. Primer pairs used to amplify major variants and other variants are depicted in grey arrows and circles (A1, Ex1, a-j, Table 1). For the variants that skip an exon, the primer spans the exon-exon junctions and is shown as a split grey arrow underneath the introns (c, g i). Protein isoforms are shown below their corresponding RNA in purple and amino acid (aa) length is listed. (B) Summary of the protein isoforms (γ) and RNA variants (var) with detailed exon/intron usage and annotations. γ2 is found primarily in adipose tissue, while γ1 is found in podocytes. Variants 1, 3, and 4 are variants of γ1 that are likely to be found in podocytes, while variants 9, 10, and 11 are additional less explored variants that would encode different isoforms. The variant 10 version of tORF4 is annotated that contains exon A1.2, but there isn’t an annotated version of tORf4 that would contain exon A1. (C) A representative gel showing the expression of several variants of PPARγ in differentiated podocytes. PCR products were generated using specific primer pairs and run on an agarose gel and stained with ethidium bromide. All PPARγ [primer pair Ex1 F, aR; 315bp]; γ2 [primer pair bF, aR; expected product 476bp]; variants 1, 9, 11 (432 bp PCR product) and Var 4 (358 bp product) [doublet primer pair, A1F, aR]; variants 3,5, and 10 (415bp product); variant 7 (341bp product); and variant 12 (503bp product) [primer pair dF, aR]. Primers are listed in Table 1.

**Table 1.**
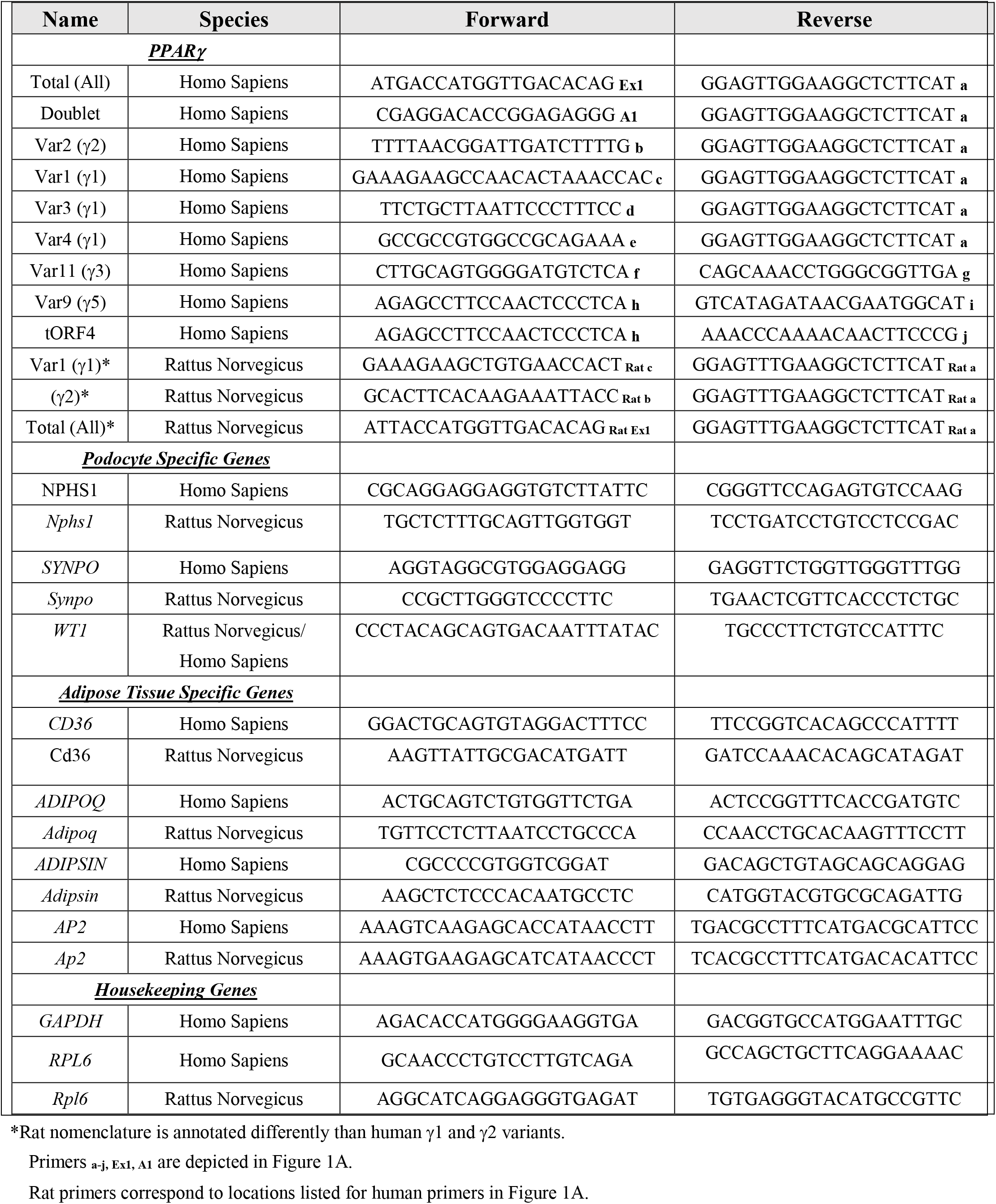
Primers Used in the Current Study

### PPAR-Promoter Responsive Element Prediction

Bulk RNASeq datasets from untreated control differentiated human immortalized podocytes (GEO GSE124622) [41] and untreated control differentiated human adipocytes (GEO GSE129153) [42] were analyzed. The average expression of control samples from the aforementioned GEO datasets were calculated and normalized expression greater than 2 were counted as detected. Genes were further cross referenced against the verified target and predicted PPAR Response Element (PPRE) genes that were downloaded from PPARgene database (http://www.ppargene.org/downloads.php) [43] and using computational genomics approach [44]. Key podocyte and adipocyte specific genes measured in this study were also queried through the PPARgene database (ppargene.org) to predict PPAR-responsive elements (PPRE) on their promoters Upon query submission, p-value and confidence level were generated if the gene was predicted as a PPAR target gene and contained putative PPREs in the 5 kb transcription start site flanking region. They were assigned into categories of confidence as: high (*p* > 0.8), medium (0.8 ≥ *p* > 0.6), and low (0.6 ≥ *p* > 0.45). A *p* value ≤ 0.45 was predicted as negative. Functional annotation of PPRE containing genes detectable exclusively in adipocytes vs. podocytes were performed using clusterProfiler (RRID:SCR_016884) [45] and plotted as many as 10 terms per cell type. The PPRE containing genes detectable in adipocyte vs podocytes the overlapping genesets, and the associated functional terms were also compared.

### Statistical Analysis

Statistical analysis was performed using GraphPad Prism Software version 8.2.0 for Windows (GraphPad Software, San Diego, CA). Data are expressed as mean ± standard error of mean and compared using unpaired Student’s t-test. P value significance is depicted as *p<0.05, **p<0.01, ****p<0.0001.

## 3. Results

### 3.1. Podocyte Specific PPARγ Splice Variants

To identify the PPARγ variants expressed in podocytes and to elucidate differences in podocytes vs adipocytes, we examined the known variants of PPARγ by reviewing the literature, scoping out the annotated versions (RefSeq database, NCBI), and by performing variant specific PCRs using custom-designed primers. A schematic of different variant forms (Figure 1A) and their exon/intron usage information and annotation (Figure 1B) is presented. Conducting PCR utilizing variant-specific primer pairs (Figure 1A and Table 1), we could identify total PPARγ expression, specific variants corresponding to γ1 (Var1 and 4) and other forms in differentiated podocytes, but not the γ2 form (Var 2) (Figure 1C), which is highly expressed in differentiated adipocytes [46]. Variant 2 (encodes γ2, a protein of 505 amino acids) includes exon B which contains a start codon. Variant 1 (encodes γ1, a protein of 475 amino acids) utilizes exons A1 and A2 instead of B in the 5’UTR of the gene, resulting in the usage of a start codon ATG located on exon 1. Variants 3 and 4 both encode γ1 as well, but they have differing 5’UTRs. Variant 3 contains an alternative exon A1.2, and variant 4 skips exon A2. Variant 11 has a similar 5’UTR to variant 1, but it skips exon 5, resulting in a frameshift and a premature stop codon in exon 6, encoding a potential product of 248 amino acids (γ3). Variant 9 utilizes a start codon located in exon 3 and skips exon 4, encoding γ5 (266 amino acids). Variant 10 or tORF4 uses a partially retained intron after exon 4 which contains a stop codon, making a product of 266 amino acids (γ6). Variant 10 version of tORF4 is annotated and contains exon A1.2, but there is no annotated variant that uses exon A1. The possibility of expression of other variants was considered using primer pairs that would amplify specific products (Figure 1C). The doublet primer pair was able to discriminate between the cumulative expression of variants 1,9 and 11 vs variant 4. Cumulative expression of variants 3, 5, 7, 10 and 12 was ruled out using primer pair for Var3 (dF and aR), which would also amplify these other mentioned variants.

### 3.2. PPARγ Splice Variants in Podocytes vs Adipose Tissue

PPARγ is a transcription factor and an established master regulator of fat cell physiology and differentiation in adipose tissue and liver [27, 47]. While podocyte specific *Pparg* KO mice illustrated its role in podocytopathies [7, 10], to understand the cell- and tissue-specific determinants of PPARγ function in glomeruli and in podocytes, we performed specific PCRs to detect the expression of total (all) variants and exclusively of variant 1 (encoding γ1) and variant 2 (encoding γ2) in human podocytes and whole kidney and rat tissues of glomeruli, whole kidney, liver and WAT. In accordance with the hypothesis of cell specific action of PPARγ, while expression of variant encoding the PPARγ2 isoform was observed in WAT, liver, and whole kidneys, γ2 variant was undetected in human podocytes and in the rat glomeruli (Figure 2A). As shown in Figure 1A, γ1 and γ2 use different, promoter sites and undergo AS. Expression analysis of several uncommon variants described in Figure 1 demonstrated that they are present at low and varying levels in the whole human kidney (Figure 2B). Most of these (variants 4, 3, 11, 9, and tORF4) were found to be expressed at very low to undetectable levels in podocytes (Figure 2B). Moreover, analysis of the variants encoding γ1 (Var 1 and 4) showed that their levels remained unchanged with PAN-induced injury of podocytes or treatment with PPARγ agonist, pioglitazone (Figure 2C).

**Figure 2.**
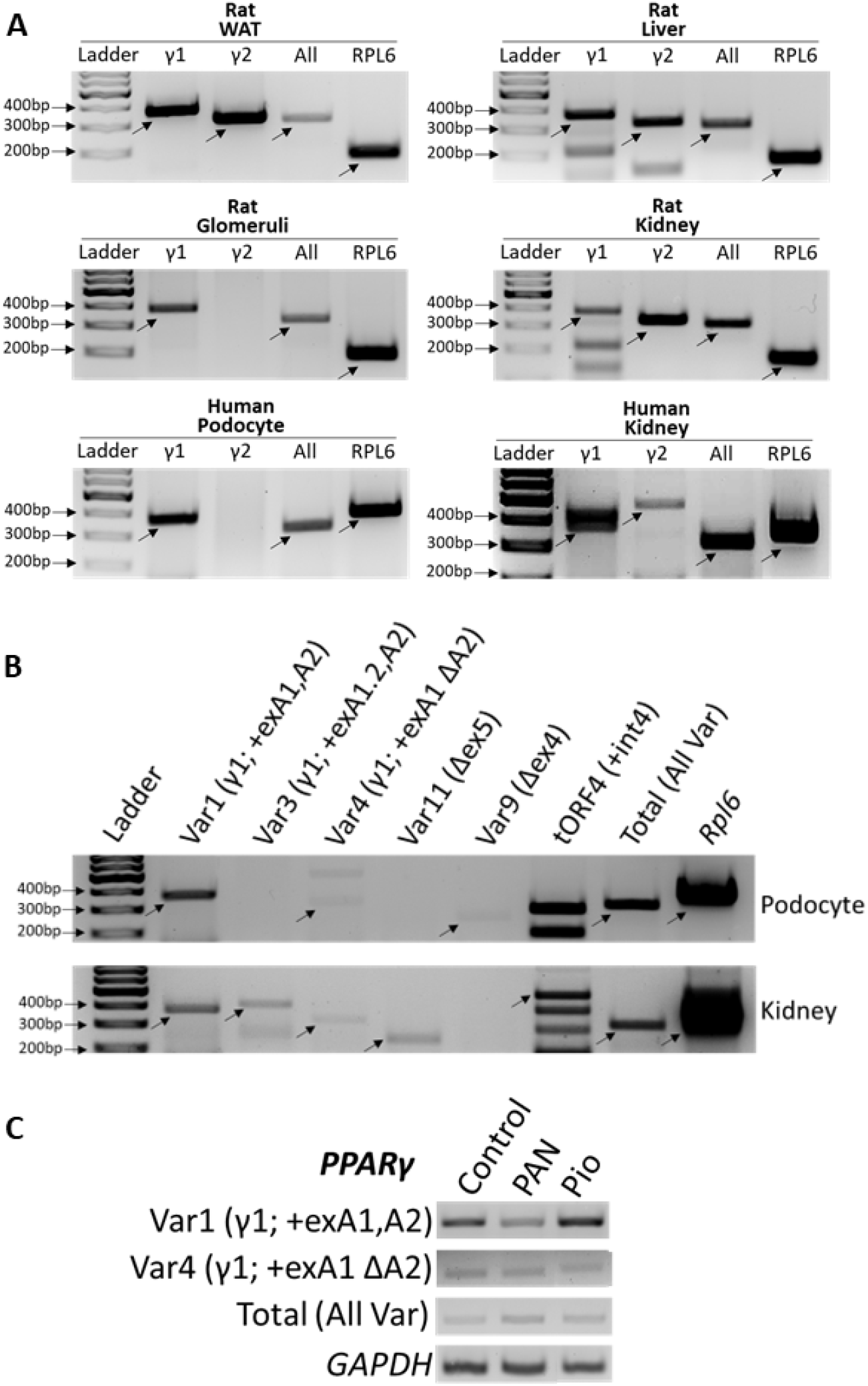
Expression of PPARγ Splice Variants in Podocytes vs Adipose Tissue. (A) Total RNA was isolated from glomeruli, liver, and WAT samples harvested from rats, and from cultured differentiated human podocytes. Expression of the RNA variants encoding PPARγ1 and γ2 (All variants), γ1 only (Var1), and γ2 only (Var 2) isoforms was measured by RT-PCR in isolated samples as well as commercially available human and rat whole kidney RNA. Rat total (all) γ [primer pair ex1 F, ex 2 R; 315bp], rat γ1 [primer pair exA2 F, ex2 R; 380bp], rat γ2 [primer pair exB F, ex2 R; 329bp], rat Rpl6 [178bp], human total (all) γ [primer pair Ex1 F, aR; 315bp], human γ1 [primer pair cF, aR; 387bp], human γ2 [primer pair bF, aR; 476bp], Human RPL6 [370bp]. (B) Several PPARγ mRNA AS variants were analyzed by RT-PCR in cultured differentiated human podocytes and commercially available human whole kidney RNA using the primer pairs drawn in Figure 1A and detailed in Supplementary Table 1. Var1 [primer pair cF, aR; 387bp], Var3 [primer pair dF, aR; 415bp], Var4 [primer pair eF, aR; 333bp], Var11 [primer pair fF, gR; 242bp], Var9 [primer pair hF, iR; 254bp], tORF4 [primer pair hF, jR; 490bp], Total [primer pair Ex1 F, aR; 315bp], RPL6 [370bp] (C) Expression of PPARγ mRNA variants 1 [primer pair cF, aR] and 4 [primer pair eF, aR], as well as total (All) PPARγ [primer pair Ex1 F, aR] and GAPDH in puromycin aminonucleoside (PAN)-injured and pioglitazone-treated podocytes. Primers are listed in Table 1.

### 3.3. PPAR-Response Element (PPRE) Containing Genes in Podocyte vs Adipocyte

PPARγ binds to its promoter response elements (PPRE) along with other transcription factors such as retinoid X receptor (RXR) or to other DNA elements in association with factors such as nuclear factor kappa B (NFκB), activator protein 1 (AP1), and in proximity to CCAAT enhancer-binding proteins (C/EBPα) [5, 24, 38]. We carried out promoter element prediction using a PPARgene database (ppargene.org) [43] and cross-referenced it with bulk RNASeq GEO datasets obtained from control podocytes (GSE124622) [41] and adipocytes (GSE129153) [42]. We found that while 1669 PPRE-containing genes were detectable in both podocyte and adipocyte datasets, 453 were found to be unique to podocytes and 77 unique to adipocytes (Figure 3Ai, Supplementary Table S1). These were further classified into subsets based on the confidence level of PPRE prediction (see Methods; Figure 3Aii). We further analyzed a select set of PPRE containing genes known to play critical roles in podocyte pathophysiology, such as *NPHS1* (nephrin), *SYNPO* (synpatopodin), and *WT1* (wilms tumor 1) or those that have established roles in adipogenesis, such as *CD36, ADIPOQ, ADIPSIN*, and *AP2* [28, 47–52] (Table 2). All the adipogenic genes depicted high level of PPRE prediction, and *NPHS1*, *SYNPO,* and *WT1* showed low to high levels of PPRE prediction. These genes which are known to play prominent roles in podocytes vs adipocytes correlated with their expression in the GEO datasets in podocytes vs adipocyte. In accordance, we also observed a marked disparity in the expression of these genes in podocytes vs white adipose tissue (WAT) (Figure 3B). While the podocyte markers (*NPHS1, SYNPO* and *WT1*) are expressed at a higher level in podocytes vs adipose tissue, expression of adipokines (*ADIPOQ* and *ADIPSIN*) and adipogenic genes (*CD36* and *AP2*) was found to be high in WAT and absent or modestly expressed in podocytes (Figure 3B, Table 2). Furthermore, functional enrichment analysis of PPRE-containing genes exclusive to podocyte or adipocyte generated distinct maps of biological processes, cellular components, molecular functions and pathways (Figure 3C-F). While the adipocyte gene set was rich in genes involved in temperature homeostasis, lipid catabolic process, lipase activity and regulation of lipolysis, podocyte gene set was rich in genes involved in cell-cell adhesion and membrane transporter activity. Adipocyte PPRE-containing gene products were found located in lipid droplet and collagen-containing extracellular matrix, and podocyte PPRE-containing gene products were found in cell-cell junction and tight junction. Of note, functional enrichment analysis of all PPRE-containing genes from podocytes and adipocytes (including overlapping genes) generated mostly overlapping maps of biological processes, cellular components, molecular functions and pathways (Supplementary Figure S1A-D). We further cross-referenced the podocyte and adipocyte bulk RNASeq GEO datasets with PPRE genes from another independent source wherein PPREs were predicted in the conserved elements of within 5000 bps of transcription site of human genes [44]. We found that out of 1074 PPRE genes, 478 were detectable in both podocyte and adipocyte datasets, 139 were found to be unique to podocytes and 22 unique to adipocytes (Supplementary Figure S2A, Supplementary Table S2). A total of 168 genes were common between the PPRE dataset and PPRE genes described by Lemay and Hwang [44] (Supplementary Figure S2B, Supplementary Table S3). A majority of these overlapping genes were detected in both podocyte and adipocyte databases (Supplementary Figure S2C).

**Figure 3.**
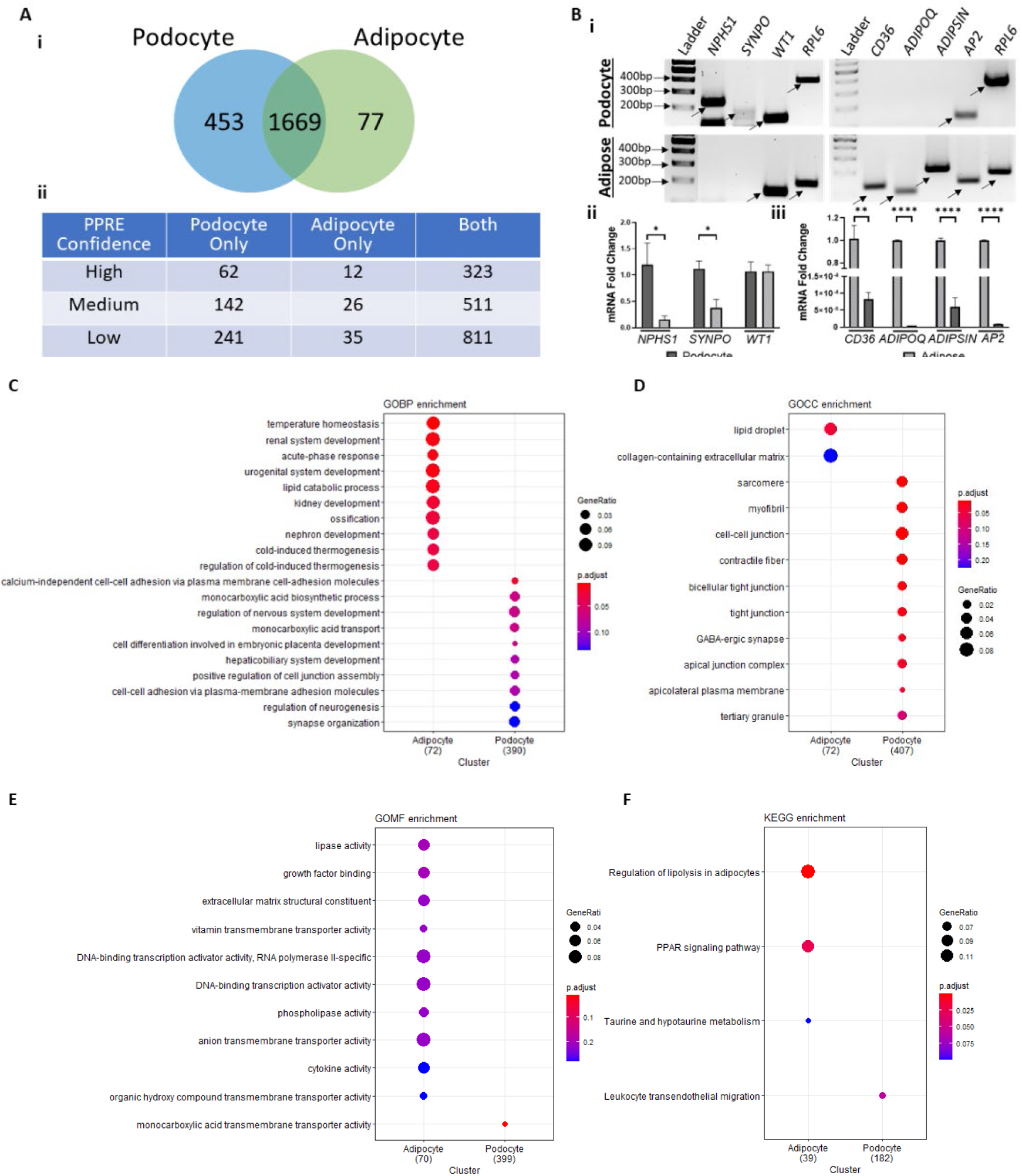
PPARγ-Response Element (PPRE) Containing Genes in Podocytes vs Adipocytes. (A) Normalized average log counts of untreated control human differentiated podocyte and adipocyte GEO datasets (GSE124622 and GSE129153, respectively) were cross-referenced against the verified target and predicted PPRE genes from the PPARgene database. (i) 1669 PPRE-containing genes were detectable in both podocyte and adipocyte datasets, 453 unique to podocytes and 77 unique to adipocytes. (ii) PPRE-containing genes classified by their confidence level. High-confidence (p > 0.8), median-confidence (0.8 ≥ p > 0.6), low-confidence category (0.6 ≥ p > 0.45). Genes with p value < 0.45 were predicted as negative. (B) Total RNA was isolated from cultured differentiated human podocytes and WAT harvested from rats. Expression of the podocyte marker genes (NPHS1 [Hu: 234bp, Rat: 138bp]], SYNPO [Hu: 182bp, Rat: 325bp], and WT1 [Hu/Rat: 134bp]) and adipose tissue genes (CD36 [Hu: 304bp, Rat: 123bp], ADIPOQ [Hu: 347bp, Rat: 95bp], ADIPSIN [Hu: 301bp, Rat: 202bp], and AP2 [Hu: 133bp, Rat: 133bp]), and housekeeping gene RPL6 [Hu: 370bp, Rat: 178bp] were analyzed. (i) Representative gels depicting the PCR products of key genes in podocytes and adipose tissue. (ii-iii) Quantitative mRNA fold -changes measured and graphically presented using RT-PCR and real time qRT-PCR quantification of key genes in podocytes and adipose tissue. Primers are listed in Table 1. (C-F) clusterProfiles generated functional enrichment of PPRE containing genes detectable exclusively in adipocytes (77) vs podocytes (453) and plotted as 10 terms per cell type for (C) biological processes, BP, (D) cellular components, CC, (E) molecular functions, MF, and (F) kyoto encycopedia of genes and genomes, KEGG. The color of the dot indicates the intensity of adj p value (smaller adj p value is more red) and size of the dot indicates the proportion of genes from the term that are present in the cell-specific PPRE containing genes.

**Table 2.** Expression of Select PPRE Containing Genes

| Gene | Cells/Tissue Tested in the Current Study | *p* Value<sup>a</sup> | Confidence Level | PPREs Predicted<sup>b</sup> | Podocyte Count log<sup>c</sup> | Adipocyte Count log<sup>d</sup> (Normalized) |
| --- | --- | --- | --- | --- | --- | --- |
| **Podocyte Prominent Genes** |  |  |  |  |  |  |
| *Nphs1* | Podocyte / WAT | 0.63696 | Medium | 4 | 2.27 | – |
| *Synpo* | Podocyte / WAT | 0.8715 | High | 5 | 7.89 | 9.03 (1.3) |
| *Wt1* | Podocyte / WAT | 0.55556 | Low | 5 | 4.91 | – |
| **Adipocyte Prominent Genes** |  |  |  |  |  |  |
| *Cd36* | Podocyte / WAT | 0.97886 | High | 3 | 3.77 | 846.26 (117.2) |
| *Adipoq* | Podocyte / WAT | 0.97153 | High | 12 | 2.59 | 112.19 (15.5) |
| *Adipsin* | Podocyte / WAT | 0.96216 | High | 6 | 4.22 | 991.32 (137.3) |
| *Ap2/Fabp4* | Podocyte / WAT | 0.99997 | High | 10 | – | 1663.6 (230.5) |
| **Housekeeping** |  |  |  |  |  |  |
| *Rpl6* | Podocyte / WAT | – | – | – | 13.42 | 96.86 (13.42) |

<sup>a</sup> Probability of PPAR target gene, higher value means a higher confidence
High-confidence ( $p > 0.8$ ), median-confidence ( $0.8 \geq p > 0.6$ ), low-confidence category ( $0.6 \geq p > 0.45$ ).
Genes with $p$ value $\leq 0.45$ were predicted as negative.
<sup>b</sup> Putative PPREs in the 5Kb transcription start site (TSS) flanking region
<sup>c</sup> Podocyte GEO Dataset # GSE124622
<sup>d</sup> Adipocyte GEO Dataset # GSE129153

In summary, this data suggests the existence of a PPRE-containing gene expression, enrichment and pathway signature that is distinct in podocytes vs adipocytes. This is likely driven by podocyte specific γ1 form of PPARγ vs the adipocyte-specific γ2 form.

## 4. Discussion

PPARγ agonism has a well-established beneficial role in the setting of diabetes, which has led to the generation of TZDs for the treatment of type II diabetes (Figure 4) [53, 54]. Its beneficial role in kidney cells beyond its favorable systemic metabolic effects in diabetes originated from numerous preclinical studies and meta-analyses [55–59]. Earlier these effects of PPARγ in influencing non-diabetic glomerular disease was thought to be through its anti-inflammatory actions on endothelial and myeloid cells [60–62]. However, recent discoveries and advances suggest the role of podocytes in mediating these effects. Seminal studies from our group and other research teams suggested that TZDs such as pioglitazone, directly protect podocytes from injury [1–5] and reduce proteinuria and glomerular injury in various animal models of glomerular disease [5–13] (Figure 4). Moreover, studies using podocyte specific *Pparg* KO mouse demonstrated that the beneficial effects of PPARγ are mediated by activation of podocyte PPARγ, thus indicating a pivotal role for PPARγ in maintaining glomerular function through preservation of podocytes [1, 5–7, 10]. However, the specific variant or isoform(s) expression of *PPARγ* and the role of its AS in podocytes and glomerular disease is unexplored. We hypothesized that AS variants of PPARγ expressed in podocytes are distinct from adipocytes. We addressed this hypothesis by performing a comprehensive analysis of different AS variants of PPARγ in human podocytes and by studying the expression of γ1 vs γ2 variant in podocyte vs adipose tissue in detail. Our findings suggest that the podocytes mainly express the y1 form, along with minimal expression of other AS forms. They do not express the γ2 form, yet they are responsive to treatments by PPARγ agonists, implicating that new selective modulators can be potentially developed that will be able to distinguish between the two forms, γ1 and γ2, thus forming a basis of novel targeted therapeutic avenues.

**Figure 4.**
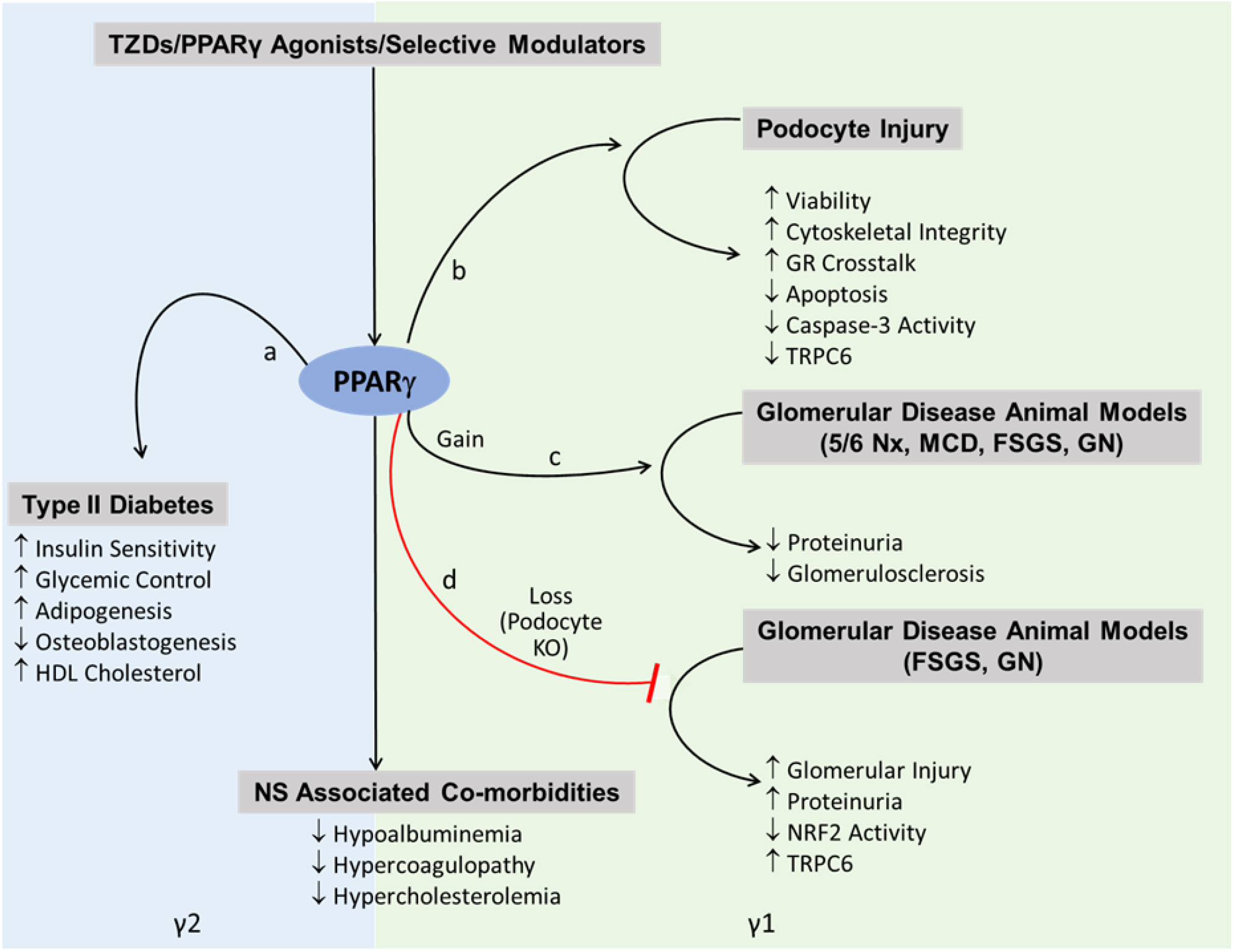
Illustration Describing the Roles of PPARγ1 and 2 Isoforms. Schematic of the suggested roles of PPARγ 1 and 2 forms in regulating podocyte/glomerular disease specific effects (right panel) vs metabolic/systemic and adipogenic effects (left panel). PPARγ agonism plays a well-established role in the setting of treatment of Type II Diabetes, and treatment with its agonists, thiazolidinediones (TZDs), leads to increased (‘a’ pathway) insulin sensitivity and glycemic control [53, 54]. However, these beneficial effects are also accompanied with adverse effects (‘a’ pathway), such as increased cholesterol, adipogenesis and decreased osteoblastogenesis [24]. TZDs or PPARγ agonists have now been demonstrated to reduce podocyte injury (‘b’ pathway) by enhancing their viability and cytoskeletal integrity and decreasing apoptosis [1–5]. A few pathways involved in this process include crosstalk with the glucocorticoid receptor (GR), decreased caspase-3 activity and decreased TRPC6 expression. Gain of PPARγ activity or its activation by TZDs (‘c’ pathway) has been shown to reduce proteinuria and glomerulosclerosis in various animal models of glomerular disease, such as 5/6 nephrectomy (Nx), minimal change disease (MCD), focal segmental glomerulosclerosis (FSGS) and glomerulonephritis (GN) [5–13, 23]. On the other hand, loss of PPARγ in podocyte specific *Pparg* KO mouse (‘d’ pathway) has demonstrated exacerbation of glomerular injury and proteinuria in animal disease models of FSGS and GN [1, 5–7, 10]. Analysis of the podocyte vs adipose tissue specific expression of genes downstream of PPARγ containing putative or identified PPREs (Figure 3) informs us that these downstream effects in podocytes/glomeruli are likely directed by the PPARγ1 splice variant, distinct from the adipocyte-regulatory γ2 variant.

Since their identification in 1990, PPARγ has been recognized as a nuclear receptor superfamily member, a ligand-dependent transcription factor and a master regulator of adipogenesis and metabolism, which accounts for the insulin sensitizing effects of its agonists or anti-diabetic drugs such as TZDs [63]. While its role in regulating adipogenesis and lipid metabolism in adipose tissue, liver and skeletal muscle is well-characterized, the knowledge of PPARγ-regulated signaling pathways in kidneys and in podocytes is scarce [5–10, 24–28, 64]. In our previous studies we speculated a cross-talk model between the PPARγ and glucocorticoid receptor and further demonstrated that while pioglitazone could provide proteinuria reducing benefits, it could also enhance the ability of low-dose steroids to reduce proteinuria [6]. This suggested the potential clinical use of these FDA-approved drugs in enabling the reduction of steroid dose, toxicity, and side effects to treat NS. We further translated these findings and demonstrated the ability of pioglitazone administration to improve clinical outcome in a child with steroid-refractory NS, which has now been extended in other NS patients as well [6, 65]. Moreover, we have recently found that targeting PPARγ with a selective modulator, GQ-16, offers an even better therapeutic strategy in NS, by reducing proteinuria and NS-associated side effects with a greater efficacy while providing reduced adipogenic effects than traditional TZDs [23]. However, to understand the pro-teinuria-reducing podocyte protective effects of PPARγ activation, it is critical to build an understanding of podocyte specific transcriptional activity of PPARγ as the lack of such knowledge is a barrier to modulating PPARγ activity effectively. The existence of two isoforms γ1 and γ2 as a result of different promoter usage and AS further complicates the PPARγ biology [29–31]. As a result, γ2 contains 30 additional amino acids at its N terminal, despite a longer γ1 mRNA sequence at its 5’UTR. We found that podocytes as well as glomeruli, express the more ubiquitous form of PPARγ, the γ1 form and not the γ2 form, which is restricted to adipose tissue and liver. Interestingly a few other spliced forms of PPARγ have been discovered in the past few years, which may confer varied functions [32–36]. Approximately 95% of mammalian genes undergo AS that can result in different protein products as well as microRNA sensitivity, mRNA stability, localization, and translation efficiency [66, 67]. Moreover, AS is associated with many cellular processes and diseases such as differentiation, cancer, and immunity to name a few [66, 68]. Moreover, expression of PPARγ1 and 2 mRNAs and isoforms in adipose tissue, liver, and skeletal muscle is known to be regulated in animal models of obesity, diabetes, and nutrition [69]. As the expression and role of the AS of PPARγ in podocyte biology is unexplored, we performed a landscape analysis of other PPARγ forms, which have been identified mostly in adipose tissues in recent years, including some with dominant negative function [32–37]. We could detect low expression of variants 4, 3, 11, and tORF4, and variant 9 was undetectable in the whole kidney. In podocytes, Var 9 was detectable at very low levels, and these additional variants were barely detectable consistently. We have recently also observed the presence of variant 11 form due to skipping of exon 5 (delta 5) in the WAT, which was particularly reduced with both pioglitazone and GQ-16 [23]. Ligand-mediated PPARγ activation has been shown to induce skipping of exon 5 which by itself lacks ligand-dependent transactivation ability, and thus acts as a dominant negative form of PPARγ [32]. In particular, recent evidence points towards distinct roles of γ1 vs γ2 variant of PPARγ governing specific and separate gene expression and metabolic functions at different stages in adipocytes [70], as well as between adipocytes and macrophages [38]. In the current study, analysis of the podocyte vs adipocyte specific expression of genes downstream of PPARγ containing putative or identified PPREs informs us that these downstream effects in podocytes are likely directed by the PPARγ1 splice variant, distinct from the adipocyte-regulatory γ2 variant (Figures 3 and 4). This is also evident from differential gene expression, enrichment and signaling pathways in the podocytes vs adipocytes. Of note, we found podocyte specific PPRE-containing genes to be functionally enriched in cell-cell junction and cell-cell adhesion, which are known to play critical roles in the maintenance of podocyte structure and function and in its normal physiology [71]. This also suggests that activation of podocyte PPARγ with agonists or selective modulators to drive these pathways is a viable therapeutic option to restore podocyte health during injury. At the same time, we acknowledge the limitations of *in silico* prediction of PPREs, species-specific differences that may exist in the location and functionality of these PPREs [72], and the involvement of other co-transcription factors that determine cell specificity of PPARγ [38]. Overall, previous evidence indicates that the: 1) activation of PPARγ results in reduction in podocyte injury and proteinuria [1, 4–6, 12, 13], 2) podocyte specific PPARγ KO results in resistance to protection of proteinuria by PPARγ agonists [7, 10], and 3) selective modulation of PPARγ enhances proteinuria reducing efficacy with reduced adipogenic potential [23]. Taken together with the findings in the current study suggesting that the podocytes do not express the γ2 form, yet they are responsive to treatments by PPARγ agonists, implicate that those new selective modulators can be potentially developed that will be able to distinguish between the two forms, γ1 and γ2, thus forming a basis of novel targeted therapeutic avenues.

PPARγ also undergoes post-translational modifications and its differential phosphorylation at serine (Ser) 273 is an important determinant of its effect on adipogenesis and insulin sensitivity [47, 73, 74]. PPARγ phosphorylation at Ser273 was demonstrated in obesity models and treatment with the PPARγ agonist dephosphorylates this residue in the adipose tissue. GQ-16, a selective modulator of PPARγ has been shown to dephosphorylate PPARγ in an *in vitro* assay like traditional TZDs [75, 76]. Since this site of phosphorylation on Ser273 is encoded by the last codon on exon 4, it remains conserved in major variants, 1 and 2, and several other variants in the current study, except for Var9, which was only detectable in podocytes at very low levels. It should be noted that the relative position is Ser243 in the variants including A1 and A2 exons instead of Ser273 in the B exon. Moreover, our recent findings suggested that relative phosphorylated PPARγ levels (at Ser273) remained unaltered during glomerular injury and with treatments with pioglitazone and GQ-16 treatments in the glomeruli [23]. These data suggest that while PPARγ-Ser273 is critical in determining the insulin sensitizing effects of PPARγ in adipose tissue, it is unlikely to be a major mechanism of regulation in podocytes or glomeruli.

In the present study, we have observed a clear indication of the disparate roles of γ1 vs γ2 in podocyte vs adipocyte function and physiology and demonstrated the presence of other AS variants of PPARγ mRNA in podocytes that would share the 5’UTR with variant 1 (encoding γ1 isoform), although the roles of these other variants in podocytopathies and podocyte differentiation is still unclear. Their detection of differential expression is a challenge as they are observed to be present at too low levels in podocytes (unlike the adipocyte variants). It might be worthwhile to pursue studies in future wherein these variants are overexpressed using reporter plasmids followed by the analysis of their effects on podocyte physiology, actin cytoskeleton and function in the presence of injury causing agents, such as PAN. The demonstration of presence of these variants also warrants their detection in kidney biopsies from patients treated with PPARγ agonists for diabetic nephropathy or type II diabetes.

In summary, we have shown that podocytes and glomeruli express the variants encoding the major form of PPARγ1 isoform and do not express the variants encoding the other major form γ2. Moreover, expression of other AS forms of PPARγ has been detected at very low levels in podocytes which may have implications in its pathophysiology. Future studies directed at developing new selective modulators that can distinguish between the two forms, γ1 and γ2, to enhance beneficial effects and reduce harmful effects of PPARγ activation has the potential to provide novel targeted therapeutic avenues.

## Supplementary Material

Supplementary Figure S1: clusterProfiles generated functional enrichment of all PPRE containing genes detectable in adipocytes and/or podocytes

Supplementary Figure S2: PPARγ-Response Element (PPRE) containing Genes in PPARgene dataset and Lemay and Hwang Manuscript detected in podocytes and/or adipocytes

Supplementary Table S1: PPRE-driven genes from the PPARgene dataset, identified in podocyte and adipocyte datasets

Supplementary Table S2: PPRE-driven genes from Lemay and Hwang manuscript, identified in podocyte and adipocyte datasets

Supplementary Table S3: PPRE-driven genes in the PPARgene dataset and Lemay and Hwang manuscript

## Author Contributions

Conceptualization, S.A.; Methodology, S.A., R.G., A.W., D.C., A.B. and C.B.; Formal Analysis, C.B., A.W. and S.A.; Investigation, C.B. and S.A.; Resources, S.A.; Data Curation, C.B., A.W. and S.A.; Writing – Original Draft Preparation, C.B. and S.A.; Writing – Review & Editing, S.A., R.G., A.W., D.C., A.B. and C.B.; Visualization, C.B. and S.A.; Supervision, S.A.; Project Administration, S.A.; Funding Acquisition, S.A.

## Data Availability Statement

The original contributions presented in the study are included in the article material, further inquiries can be directed to the corresponding author.

## Acknowledgments

The authors thank Dr. Jeffrey Kopp at the National Institute of Health and Dr. Moin Saleem at the University of Bristol for sharing the immortalized human podocyte cell line, and members of the animal resource core at the Abigail Wexner Research Institute, Nationwide Children’s Hospital (NCH).

