## Supplementary Material for "Alternatively Spliced Landscape of PPARγ mRNA in Podocytes is Distinct from Adipose Tissue"

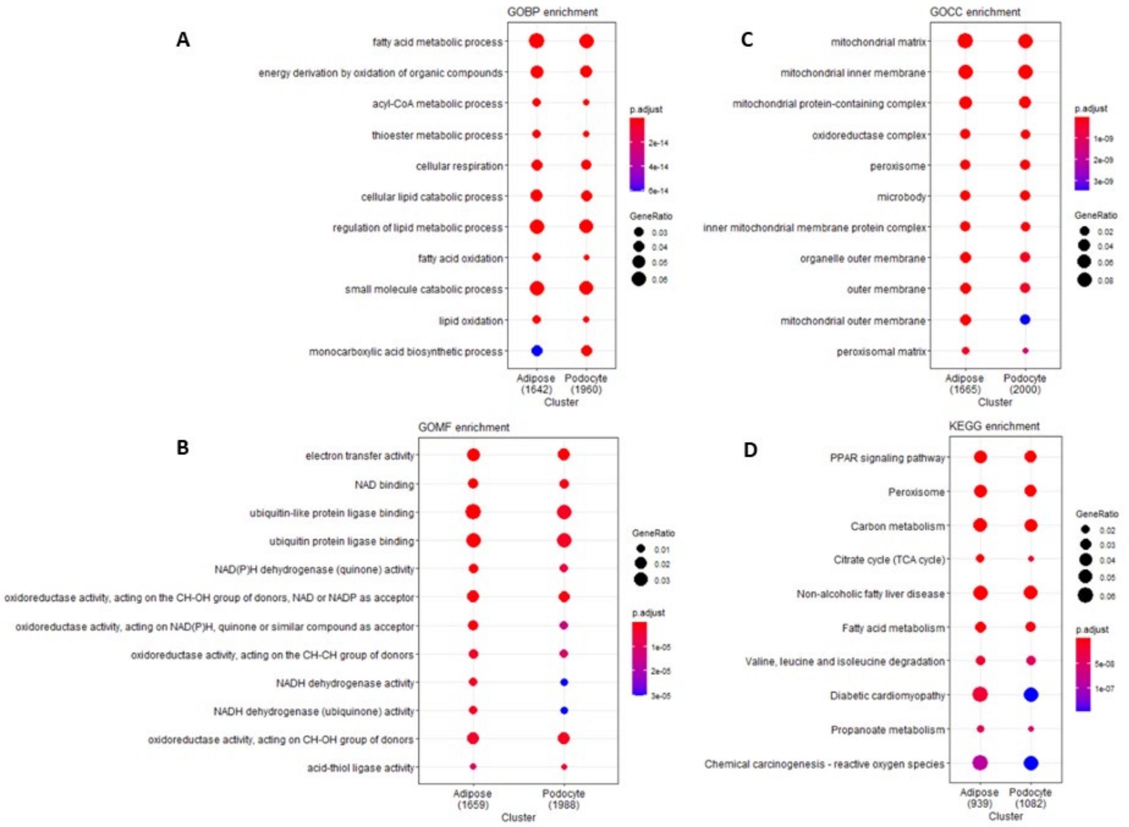


**Supplementary Figure S1.** ClusterProfiles generated functional enrichment of all PPRE containing genes detectable in adipocytes (1776) or podocytes (2122) and plotted as 10 terms per cell type for (A) biological processes, BP, (B) cellular components, CC, (C) molecular functions, MF, and (D) kyoto encycopedia of genes and genomes, KEGG. The color of the dot indicates the intensity of adj p value (smaller adj p value is more red) and size of the dot indicates the proportion of genes from the term that are present in the cell-specific PPRE containing genes.

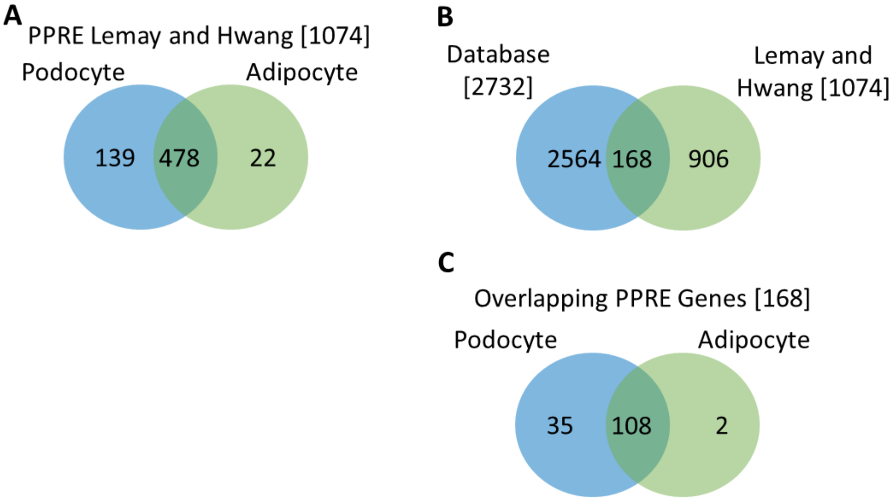


**Supplementary Figure S2.** PPARγ-Response Element (PPRE) Containing Genes (A) in Podocytes vs Adipocytes. Normalized average log counts of untreated control human differentiated podocyte and adipocyte GEO datasets (GSE124622 and GSE129153, respectively) were cross-referenced against the human PPRE predicted genes described in Lemay and Hwang manuscript. 478 PPRE-containing genes were detectable in both podocyte and adipocyte datasets, 139 were unique to podocytes and 22 unique to adipocytes. (B) PPRE-containing genes from the PPARgene database and Lemay and Hwang. (C) Majority of the overlapping PPRE genes (168) between the PPARgene dataset and Lemay and Hwang manuscript were detected in podocytes and/or adipocytes.

**Supplementary Table S1: PPRE-Driven Genes from the PPARgene Dataset, Identified in Podocyte and Adipocyte Datasets**

| Podocyte Not Adipocyte (453) | Podocyte And Adipocyte (1669) | Adipocyte Not Podocyte (77) |
|---|---|---|
| KRTCAP2 | AACS | WISP2 |
| PYROXD2 | ELMO2 | A2M |
| XDH | BTG2 | SLC38A4 |
| SLC4A1 | NDUFS6 | PBX1 |
| PBX4 | TACC2 | HP |
| TMEM139 | PDHA1 | PROX1 |
| SYCP3 | ABCB7 | SIX2 |
| GDA | HSPB1 | S100B |
| SLC15A2 | PFKP | MGP |
| MREG | ARRB2 | LGALS12 |
| PHYHIP | DECR1 | CILP2 |
| UNC5A | CMTM8 | PDE3A |
| LRRN1 | PRADC1 | AREG |
| MRPL38 | CSPG4 | ARHGAP20 |
| TMEM184A | IGF1 | GREM2 |
| CCL22 | ITPK1 | CD302 |
| LRFN5 | G3BP1 | ISLR |
| SLCO2B1 | HSD17B10 | SRPX2 |
| ZMYND15 | LMAN1 | DIO2 |
| A1CF | UQCR11 | IL1RN |
| CLDN16 | GDF15 | MAFB |
| ISL1 | CYCS | SLC24A3 |
| C1QTNF3 | GPSM2 | ALPL |
| HSPA1L | HIC2 | PTGER3 |
| STARD5 | GYS1 | XPNPEP2 |
| ITGB1BP2 | DTNBP1 | HSPB6 |
| CORO2A | TAF9B | TC2N |
| GMPR | TRIB3 | CILP |
| FRRS1 | REEP6 | TUSC5 |
| SHBG | CLUH | GPX7 |
| SLC16A5 | CDK14 | AADAC |
| LLGL2 | NDUFS1 | MRVI1 |
| FZD9 | SOD2 | PLA2G2A |
| KL | AES | CHST1 |
| ATP1B4 | ARL2 | SLCO3A1 |
| CDS1 | KCNK3 | CITED1 |
| ADAM19 | NDUFAF4 | CCL7 |
| ADAM32 | STK16 | OLFM1 |
| FNDC8 | NDUFAB1 | CHKB |
| CYP1A1 | FGFR1 | SLC19A3 |
| ERO1L | AGPAT1 | RORB |
| XRCC6BP1 | CERK | RAMP1 |
| KCNQ5 | PDLIM2 | ZBTB16 |
| NPHS1 | ATP5D | ASPA |
| SLC39A5 | RAB40B | EDNRB |
| ADRA1A | RCL1 | PCK1 |
| NRG4 | PIR | CIDEC |
| FAM154B | LONP1 | EBF1 |
| SCN11A | PPCS | HTRA3 |
| CDH24 | MRPL1 | TRIM69 |
| LCA5L | SYNGAP1 | MTERF4 |
| EML5 | ARL6IP1 | THRSP |
| FAM195A | BET1L | FMO1 |
| SETD8 | FIBCD1 | RNF5 |
| NR1I3 | PTCD3 | ANGPT1 |
| ACAD11 | FADS1 | GLMP |
| TJP3 | PCYT2 | PLIN1 |
| CHRM4 | SERPINF1 | CD163 |
| NIPAL4 | TSC22D1 | MYH14 |
| TMEM25 | ATXN10 | GADD45G |
| RRAD | ECH1 | RBP4 |
| SNAI1 | FAHD1 | CDO1 |
| SLC35F1 | NMNAT1 | SEPP1 |
| PFKFB1 | RHOT1 | PLCL1 |
| CAMK2B | ELOVL5 | CACNA1A |

| | | |
|---|---|---|
| NGFR | EPS8 | PRR15 |
| GRHL1 | DIP2A | PLBD1 |
| CCDC132 | SHC1 | RASD1 |
| ANO9 | ACTR8 | SPON2 |
| IL1A | BRCA1 | APCDD1 |
| UPK3B | GEN1 | KLF15 |
| TMEM79 | FAM96B | FABP4 |
| KRT8 | S1PR2 | HSD11B1 |
| SCN4B | PAPSS2 | LAMA4 |
| GATSL2 | ACSL3 | SDCBP2 |
| MYCL | GRPEL1 | PARM1 |
| GLI1 | USP6NL | AQP7 |
| TRABD2B | ROCK2 | |
| RAB11FIP4 | PPP1R15B | |
| SLC25A34 | HES1 | |
| SLMO2 | ALDOC | |
| IL21R | VGLL3 | |
| TLR5 | SEL1L3 | |
| FAM78A | PDCD4 | |
| CD200 | CEP350 | |
| INCA1 | FAM73A | |
| SGK2 | RHOC | |
| KLRG1 | PPP3CA | |
| NEURL3 | DDB2 | |
| LAMC3 | NEIL1 | |
| TOX | TSPAN13 | |
| CHDH | FITM2 | |
| CLDN10 | SPIRE2 | |
| PTP4A3 | MKNK1 | |
| ROPN1L | DNAJC15 | |
| ERC2 | RAD23B | |
| RBP7 | MGAT4B | |
| HSD17B7 | NCOR2 | |
| GAL3ST1 | IL17RC | |
| TRIM14 | LRRC8D | |
| AIF1L | ALDH18A1 | |
| NRSN1 | CEP170 | |
| SHANK3 | UFM1 | |
| ABHD16B | MAOA | |
| PFN4 | TPI1 | |
| SPNS1 | APOE | |
| SLC23A3 | ALDH2 | |
| GPR160 | EMP2 | |
| SLC14A2 | IP6K1 | |
| LSMEM2 | SRSF6 | |
| HUNK | MRAP | |
| GPT | QDPR | |
| GLTPD2 | HSPD1 | |
| LRRC39 | EIF1 | |
| GYLTL1B | RIN2 | |
| REEP1 | TMEM98 | |
| UBE2T | CDK1 | |
| PRELID2 | SLC38A10 | |
| PROCA1 | OPHN1 | |
| TRPV1 | NDUFA8 | |
| FAM84B | FABP5 | |
| FGFBP1 | GCC2 | |
| FOXG1 | GLTSCR2 | |
| FMNL1 | TOR3A | |
| CDK18 | PLIN2 | |
| MCF2L | CD82 | |
| PDZK1IP1 | CHMP1B | |
| CHGB | PPM1K | |
| TMCO6 | PMM1 | |
| DBP | GCLM | |
| UCKL1 | CPEB3 | |
| STARD8 | BCL2 | |
| PPP2R4 | NFKB2 | |
| CCDC103 | PCDH7 | |
| CSDC2 | ADI1 | |

| | |
|---|---|
| EMB | SCP2 |
| KRT80 | ALCAM |
| ANKMY1 | SH2B2 |
| HHEX | ID1 |
| ACY1 | COL18A1 |
| ALOX5AP | ERLIN2 |
| ADHFE1 | PPP1R9A |
| GNAZ | PDZRN3 |
| PSD | TIMM22 |
| CLDN3 | GPD1 |
| BTN2A2 | PTRF |
| RASGEF1B | PRPS2 |
| ACSBG1 | MMP15 |
| POLR2F | ALAD |
| AQP3 | ASS1 |
| STAB1 | GSTM4 |
| ASPDH | RGS4 |
| CLSTN3 | UQCRC1 |
| NFE2L3 | SNX6 |
| SH3D21 | PPP2R5A |
| CD74 | MRPL34 |
| UBXN11 | SAMM50 |
| GPR35 | SRP72 |
| BCKDHA | ESRRA |
| PLEKHH1 | LDHB |
| PIK3R2 | LRRC8A |
| SMPD3 | DDX49 |
| FOXS1 | GNA11 |
| GLDC | TLCD2 |
| KRT18 | EBPL |
| PTPN6 | RHBDF1 |
| INHBE | SNX25 |
| CCDC85C | KCNE3 |
| HEY1 | CPEB1 |
| ITGAX | ACOT7 |
| PLIN5 | ENDOG |
| ADORA2B | EZR |
| PHEX | ABHD12 |
| PCDH17 | PTGES |
| TOMM70A | PEX16 |
| KRT19 | FGF1 |
| GIPC3 | WDR6 |
| NEURL4 | FUT4 |
| ADAP2 | DNAJA3 |
| NUDT13 | ITFG2 |
| DAK | EPHA4 |
| RARRES1 | TMEM11 |
| SIM2 | IMMT |
| LRG1 | UBE2L6 |
| NRXN3 | AGGF1 |
| RHPN2 | SPRY2 |
| SEMA4A | TPBG |
| POU5F1 | LZTFL1 |
| VSIG2 | GSTP1 |
| CLDN1 | GMFB |
| SLC27A2 | LTBR |
| DDO | AGTPBP1 |
| NHLH1 | PRR5 |
| TRIM46 | TLE2 |
| LCAT | RPRD1B |
| ARHGEF25 | STAT5A |
| ST6GAL1 | FKBP4 |
| TSSK2 | PNPLA6 |
| PSMA6 | GIPR |
| CEACAM1 | PERP |
| UCP3 | YIPF2 |
| COL8A2 | FHL1 |
| CYP8B1 | TNFSF12 |
| LIN28A | PSMD5 |
| DHX58 | FAM126B |

| | |
|---|---|
| GCN1L1 | LACTB2 |
| AMOT | GANC |
| TIMD4 | RBPMS |
| ADAM11 | ACO2 |
| CDA | TPMT |
| TEX11 | ARID5A |
| GAREML | LOXL1 |
| GJB4 | NFE2L2 |
| FBXO2 | KLF4 |
| AMIGO2 | ACSF2 |
| TMEM26 | PFKL |
| EVPL | SNN |
| NR1D1 | TMEM140 |
| CACNA1G | GABBR1 |
| AS3MT | POLD2 |
| HYAL1 | LSM3 |
| BEAN1 | ACOT2 |
| GNB3 | PPL |
| L3MBTL1 | PPARGC1A |
| AGPAT9 | ETFA |
| E2F8 | AHRR |
| PVRL1 | RASSF4 |
| CHRNE | MTHFD1 |
| EHF | SOX9 |
| ATXN7L2 | CASC4 |
| ACOT1 | SYTL4 |
| CPSF4L | CEBPD |
| RAD51B | VRK2 |
| FBXL6 | JUND |
| NGF | MOSPD1 |
| ALOX5 | PHB2 |
| MDGA2 | EHBP1L1 |
| FGFBP3 | NDRG2 |
| NXPH3 | COG4 |
| TGM2 | NDUFB8 |
| DEFB1 | DIXDC1 |
| YBX2 | DNAJC11 |
| ARHGAP30 | TIMM17A |
| ARPP21 | CADPS2 |
| CIB2 | SEMA3G |
| DSCAML1 | EEPD1 |
| PRICKLE3 | AMMECR1 |
| NEURL2 | LRRC2 |
| MEOX1 | DLST |
| SYT10 | NEDD4L |
| DOCK8 | NCAPD3 |
| PLEKHB1 | FST |
| GRB7 | LDLR |
| BDNF | SCRN1 |
| RASAL1 | PROSER2 |
| SEMA3E | COQ5 |
| RASD2 | PMVK |
| TMEM143 | SGK1 |
| LRRC17 | DDI2 |
| ELF3 | RNF7 |
| CXCR2 | POLR3G |
| PODXL2 | CPD |
| CELF6 | FBF1 |
| ADCY5 | IRF7 |
| ESYT3 | PELI3 |
| DEGS2 | TBC1D8 |
| MERTK | HSPA2 |
| ISL2 | SLC38A9 |
| CCRN4L | ACAP3 |
| TNNC1 | NFKBIZ |
| TRIM55 | RAB20 |
| STBD1 | PPOX |
| FOXO6 | TAF9 |
| CLDN7 | DGAT1 |
| FAM69B | METTL5 |

| | |
|---|---|
| CTF1 | ACADVL |
| JMJD7 | ISCU |
| HDGFRP3 | RAB15 |
| SV2B | PHGDH |
| WT1 | FDX1 |
| C2CD4C | XIAP |
| NCCRP1 | WNT5A |
| CLDN23 | SH3BGRL2 |
| MYH3 | DOPEY2 |
| RSPH1 | PEX26 |
| PRUNE | PTGS2 |
| ITGB6 | SLC29A3 |
| SLC1A2 | SDHD |
| RGL3 | IMPAD1 |
| CERS1 | ADAMTSL4 |
| MAGIX | OPN3 |
| ITPKA | SQLE |
| RHBDL3 | DAGLB |
| CD27 | HCFC1R1 |
| RHEBL1 | GLIPR2 |
| SPIN4 | WBP11 |
| SULT2A1 | LYPD3 |
| SOX17 | TEX2 |
| GARNL3 | MAP1A |
| SLC6A13 | MAP4K3 |
| TINAG | COX7B |
| RUNDC3B | TMEM185B |
| IL17RB | ANKRD17 |
| RNFT2 | GRPEL2 |
| USP2 | SNTA1 |
| ABCG4 | H2AFY2 |
| PLCD4 | FOXN2 |
| COQ3 | NT5E |
| LACE1 | FAM129B |
| ADRB2 | ITGA6 |
| TRERF1 | NCK2 |
| TUBB3 | SLC16A2 |
| S100A1 | TRIM35 |
| SH3TC1 | SC5D |
| GRB14 | USP11 |
| NTRK3 | IMPA2 |
| MMEL1 | PRKDC |
| CHODL | SULF2 |
| IL12A | NET1 |
| GDPD1 | MVD |
| PUS10 | ACLY |
| KRT81 | DNAJC1 |
| POPDC2 | ARHGAP12 |
| FAM26F | CCNDBP1 |
| ASGR1 | MEX3D |
| PVRL2 | FKBP5 |
| IL6 | SCCPDH |
| SNAI3 | GPX4 |
| CHAC1 | CCDC85B |
| EPOR | PRMT1 |
| ANGPT2 | FLI1 |
| SSPO | TMOD2 |
| ITGAM | FAM107B |
| CCNO | IRS2 |
| RGS5 | PPDPF |
| ABCG2 | ALG9 |
| ACRV1 | PIM3 |
| HPD | ANKRD40 |
| HNF1A | COMTD1 |
| TMEM45B | ACOT9 |
| STXBP2 | SPATS2L |
| OSCAR | ADCK2 |
| DNMT3L | SLC48A1 |
| PLCG2 | UQCRQ |
| BCMO1 | RIMS1 |

| | |
|---|---|
| ITGA2B | RCN1 |
| CYP1A2 | KAT5 |
| CABP4 | BCAR3 |
| SRXN1 | MAP3K6 |
| EPN3 | DHCR7 |
| STARD10 | TPD52L1 |
| HOMER2 | CRAT |
| ANAPC2 | HN1 |
| KCNS1 | POR |
| INA | VEGFA |
| SLC16A6 | CSRNP1 |
| SUV39H1 | PAK1IP1 |
| LY75 | TRUB1 |
| PCSK4 | ELL |
| KCNAB2 | FAM89A |
| DMBT1 | VAT1 |
| SPNS2 | SENP3 |
| FOXA1 | ACKR2 |
| VNN1 | EEF2K |
| SLC16A13 | TUFM |
| TEX22 | MPDU1 |
| ALDH3A1 | MYC |
| GALNTL6 | DECR2 |
| IGDCC4 | SPTLC3 |
| KCTD6 | MUS81 |
| DDN | PHYH |
| ARAP2 | ACAA2 |
| TMEM35 | TMEM53 |
| ABHD3 | DAG1 |
| PNMAL1 | TIMP1 |
| ANKRD2 | RUVBL2 |
| RAD54L | PLOD2 |
| TTYH2 | ISCA1 |
| ABCB4 | UCK1 |
| GPR146 | MRPL9 |
| SORD | RUNDC1 |
| MFSD4 | SRSF3 |
| FRK | ICAM2 |
| PPAP2B | LFNG |
| HOXA4 | AUH |
| TGFA | SLC25A39 |
| MATN1 | GNA13 |
| RTP4 | MBD2 |
| NPPC | TNFAIP3 |
| COX6B2 | PSRC1 |
| THOC6 | ALAS1 |
| EFTUD1 | FOXO1 |
| ACBD4 | MRPL37 |
| AQP1 | NDUFV2 |
| ACER2 | PDK4 |
| CYP26B1 | DUSP10 |
| CNKSR1 | FARS2 |
| DNASE1L2 | CCDC28A |
| ATAT1 | PKP2 |
| TMEM128 | AMPD3 |
| TMOD1 | PIAS1 |
| RSAD2 | TMEM134 |
| PLLP | STAT1 |
| GLRB | GCLC |
| AMN | ZC3H14 |
| MSLN | COL8A1 |
| LMTK3 | COLEC12 |
| SLITRK1 | RPP30 |
| SLC45A3 | BLVRB |
| MEST | GTPBP2 |
| PDZK1 | CWC25 |
| RNMTL1 | ELK4 |
| RTKN | MKKS |
| APOA1BP | BANF1 |
| GPRIN3 | FAM13A |

| | |
|---|---|
| CRTAC1 | SLC22A3 |
| CDCP1 | BSG |
| ERCC6L | GADD45B |
| EDN2 | SLC26A6 |
| SHISA2 | BBX |
| CCDC121 | UCP2 |
| CGN | DTX4 |
| TLCD1 | BNC1 |
| LTC4S | ALS2 |
| PRSS8 | PABPN1 |
| WNT4 | NDUFB3 |
| PPFIA4 | SLC22A5 |
| PAQR9 | SLC25A42 |
| KIF21B | FKBP8 |
| KCNJ5 | PTCD2 |
| TSNAXIP1 | MMP11 |
| DLX1 | MIPEP |
| DOK7 | UCHL3 |
| CSF1R | PPARD |
| RBP1 | ANXA2 |
| MYOM3 | SPEG |
| DCAF12L1 | TBC1D24 |
| PSMB10 | STK10 |
| PAM16 | ABCD3 |
| GATM | HDDC2 |
| RUNDC3A | INPP5F |
| SPINT2 | VPS26B |
| RNF128 | MYBL1 |
| CD274 | PAQR7 |
| DAPK1 | INPPL1 |
| VAV3 | FAM73B |
| TRPC3 | CDC27 |
| HCAR1 | MDGA1 |
| MNX1 | COPS6 |
| SERHL | BRI3BP |
| ADRBK2 | CNN2 |
| TNNT1 | GPC4 |
| LIPT2 | LMO7 |
| | SOGA1 |
| | COL4A2 |
| | BPGM |
| | TCF7L1 |
| | SLC12A7 |
| | HMOX1 |
| | CITED2 |
| | DTX2 |
| | BMF |
| | FAM126A |
| | SIAH2 |
| | PFAS |
| | ABHD5 |
| | MKNK2 |
| | TMBIM1 |
| | EFNA5 |
| | FCGRT |
| | NXN |
| | IGF2BP2 |
| | URGCP |
| | PNPLA3 |
| | MAF1 |
| | ACADM |
| | GIPC2 |
| | SLC22A23 |
| | DUSP3 |
| | ANGPTL4 |
| | MPZ |
| | NAMPT |
| | HECTD1 |
| | SDSL |
| | SLC31A1 |

HACL1
RAD23A
FAM102A
SFXN1
RNASEH2C
IRAK2
GHITM
NDUFB7
FOXN3
TXLNG
MFGE8
YAF2
SP110
SH3BP2
PXMP4
TFB1M
DCAF4
ATP9A
LCLAT1
TSKU
ETV3
PDLIM1
UQCR10
CLIC4
TLE3
DDX31
PPP1R14B
LRP1
MESDC1
MMP14
SERPINI1
HDHD3
PLAT
NUPL2
TDRD3
PPP2R5B
KLHDC3
GNAI1
SLC25A4
CYB5B
NDRG1
SESN1
TMED8
SDHAF2
ARHGAP1
GBE1
PIP4K2A
SOD1
AFAP1
PACS1
FSCN1
TMEM136
SIRT5
MSRB2
ZFP36L1
CBR3
NR4A1
WDR74
ST6GALNAC4
MCM7
ATP5G3
METTL21B
IGFBP6
ABCB9
TSC22D2
CFL2
RASA2
SLC22A15
STX18
DBI

BTBD1
LPIN2
ZHX2
PDP2
SMARCA4
ARMC9
RPS6KA4
MRPL14
C1QTNF6
TTC8
AGL
CEP55
PALMD
HS6ST1
TTC39B
ENPEP
TIMM23
OAS2
DEGS1
FAIM
LIPG
TBL1XR1
MGST2
SLC25A11
STARD4
OSBPL1A
RPL3
KLB
ACSL1
TTI1
TNFRSF21
CDC42EP5
SLC6A8
CDC45
TMEM38B
TAZ
NFE2L1
STIP1
NT5DC2
CHST7
CCT3
SLC25A20
HSPE1
IER5
MAPK13
EIF4EBP1
APLN
C2CD2L
SNRK
GUSB
PMPCB
NID2
HYLS1
CHEK2
NUDT16L1
PRKRIP1
PI4K2B
MCM4
TCEA1
RNF111
EGLN2
CHST12
CLPX
MRPS36
RAB34
PJA1
CHMP1A
PHLDA2
BZW2
HADHB

BAHD1
TST
LYSMD3
ERLIN1
GPX8
CHCHD10
TNIP1
GLRX
RHOG
CYB5R3
CDK2
GPAM
CDC37
FBXL7
LRRC8B
SEC14L2
POLR2A
MBNL3
YIF1A
NR2F6
TMEM218
GRINA
TFPI
H2AFV
TMEM87A
TBC1D22A
SMTN
HIP1R
DGKA
PQLC1
ENO1
SBDS
DET1
GALT
AZIN1
SLC7A7
RNPS1
MB21D2
ISOC1
NR2C1
SLC25A10
TMEM64
ARG2
MKI67
PEX5
LENG1
ZFAND5
MAFG
MAP1LC3A
TOMM5
SLC41A1
ZCCHC2
PAXBP1
LOX
FIZ1
YARS
ELOVL1
UBE2E2
LAMB2
DTD2
PBXIP1
IMMP2L
MARCKSL1
RAB11FIP5
BAG4
CDKN1A
CHCHD6
LPL
CAT
CUL7

CPOX
SPSB1
RNF10
TPM2
PPP1R11
TMEM120A
WDR48
PIK3R1
ACACB
HOXC11
MAF
TRAK1
CSTB
ADM
HILPDA
PREPL
STAMBPL1
PTGR2
CRYL1
ACP6
SELM
PPP1R1A
ETFB
HADHA
IDH1
XRCC5
PLA2G16
RBM39
HDAC10
ELMOD3
ZFYVE27
RGS17
RUFY3
PCTP
FASTK
ABCC1
MRPL51
TMEM135
HAX1
KDM2B
NEK9
SLC25A45
CTSA
TOMM40
UBFD1
MMADHC
MBOAT7
HOXC8
AIG1
APBA3
APEX2
DEDD2
HAUS7
HMGCS1
ZBTB3
MRPL23
ITGA7
ORC4
CNOT3
MTIF2
HIBADH
TXN2
TSPAN4
MPRIP
ZFAND2A
CPT1A
SREBF1
EFR3A
SUCLG2
EPAS1

AMPD2
IFNAR2
LPIN1
SLIT2
CCP110
ITGAE
GAPDH
ELK3
COMMD6
SLC39A1
NARS2
PSME2
LIMK1
FAM20C
TSPAN17
MMP19
HOXB7
TMTC1
GSTM2
PCOLCE2
ATP6V0E2
PEAR1
ME1
PGS1
KDSR
MCM8
SETD4
SLC4A4
RHOBTB1
ACOT8
BHLHE41
SMAP2
FAM101B
PEBP1
MRPS18B
AOX1
RPL14
PCSK7
RMND1
LTBP3
ABTB1
CTTN
KCNJ8
PASK
SLC16A1
EHHADH
RANBP9
USP8
TMEM147
LNX1
INSIG1
CFD
SYNGR1
APOC1
TMEM37
ERC1
WDR73
YPEL3
MLYCD
SCO1
NDUFA5
ZBTB40
SDHA
NCLN
TMEM206
SAT1
PDLIM5
BRWD1
THNSL2
RCBTB2

C1QTNF1
VEGFB
LETMD1
UQCRC2
PVR
ISCA2
S1PR3
ANKZF1
TMTC2
UCK2
LBR
PGK1
WLS
CDC25A
KLF3
ANKRD37
NAT14
RCN3
RRN3
B3GNT5
TEX264
COBLL1
CHPT1
RPTOR
ERGIC1
EIF4EBP2
MEF2D
IER2
WASL
MACF1
TRAF3IP2
MDFIC
MED20
ADD3
RAP1A
ERI1
TRIO
ROGDI
MRPS35
RASSF3
RDH13
HOXC13
SLC25A37
HMGB3
CTNNBL1
MID1IP1
ECHDC1
NDUFA6
KLF11
NDUFAF3
PHLDA3
DSP
GALK1
NUP214
IRF1
DPYSL2
SOX13
SSH2
SDHB
SOX4
TSPAN3
DNAJB1
AK3
RHOF
USP20
NRP1
NR2F1
PLIN4
JAM3
FASN

MRPS14
PHLPP2
LAPTM4A
PRDX6
GPNMB
TTF2
SLC25A26
DOLK
ACVRL1
ACYP2
PSMA1
NFATC4
TMEM60
NR1H3
ABCG1
RGS3
SQSTM1
SNRNP70
SLC35F6
UAP1L1
ACAT1
PLK3
SMAD3
RALY
PHKA2
ATF3
UCHL1
FBXO22
DGKH
ELOVL3
TAGLN2
SLC16A7
NDUFA11
PUSL1
RASL10B
MRPL19
TAPBPL
PRKAR2A
KLHL23
PEX14
CNOT4
KLF16
PGRMC2
PLTP
THOC2
LGALS1
MINK1
DNAJA4
SLC25A1
EZH2
EXOC6B
SRI
SOCS6
SLC25A5
MYO1C
YWHAE
WDR1
AFG3L2
ADAT2
CNKSR3
MGST3
HBEGF
CDKN1B
R3HDM4
NDUFS8
ZC3H3
ABCA1
DDAH1
MGAT1
TNFSF10

TIPARP
CRYAB
CHEK1
MRPL47
MUC1
SLC40A1
RNF34
ACOX1
CD36
SNHG11
CENPH
ANAPC13
HSD17B11
SEC31B
IDH2
HSD17B4
ZNHIT1
SLC12A6
EFHD1
COPZ2
CPSF3L
NUDT7
NOP2
NNAT
ADAMTS9
GPCPD1
BCL9
MRPL28
STIM1
PIGV
CDK2AP2
LETM1
APBB1
CAMK2D
ALG14
LIPE
SLC25A44
SFSWAP
FGD4
CLP1
PTGES3
KIF16B
CASP8
CS
BNIP2
NRP2
CPSF7
MED10
PTRH1
CDH2
COX17
GTF3A
ERMP1
MRPL54
SIPA1L2
CCNG2
RCAN2
EMG1
WDR5B
NCEH1
PPARG
HIRA
SLC47A1
ORMDL3
CORO2B
PAN2
DMPK
TECR
RNF144A
CBR4

SLC1A5
LPXN
PUS1
SMYD5
PEX1
TUSC2
MYL6
FADS2
HSPA4L
ENAH
SMAGP
PARP14
PTGFR
TRIM37
ABHD15
CRY1
ABCC4
PDRG1
FAR2
ZCCHC24
PIM1
THBD
DNAJC7
PARP11
NOL3
IFITM2
ACTR6
MRPS18A
LYRM5
CHCHD7
SERINC2
PPP1CC
TM7SF2
CDCA7L
LEPR
CCDC109B
MTOR
LTBP2
SRP68
NDUFS2
SLC2A8
SLC25A22
ILK
STK38L
MRPS7
AOC3
MTHFD2
FABP3
SLC27A1
TRPV2
RALGAPA2
CAMKK1
OAS3
ANXA6
LRIG3
VAPB
AP2A1
FZD7
AFF4
MRPS33
STOM
AIM1
SMARCA5
GAS2L3
UNC79
PDE8A
PLA2G4A
SMOX
ARPP19
FKBP10

SMAD7
VLDLR
VEGFC
DBT
SDC4
HIGD1A
CALR
WARS
NUDT6
ABCC3
NAA50
PIP5K1A
CARS2
BTBD2
MMACHC
BEST1
REV3L
TWF2
CYC1
PEX13
DISP1
SGTB
CHUK
ZDHHC12
DENND4B
PDK2
SRGAP2
SLC38A2
SYNJ2
TRIM26
ST3GAL6
ATN1
TRPT1
NUDT14
CAP1
DUSP1
SLC44A1
GFM1
NPR3
KBTBD11
PRPS1
NDUFB5
LIPA
SETD6
CDC42EP2
OXR1
GAMT
MRC2
MIDN
FRAT2
NAA10
BAD
LRPPRC
NDUFAF7
MRPL35
ACO1
RBX1
GRHPR
MYO18A
MEGF9
GABARAPL1
SLC4A2
ARRB1
PFKFB3
SCAMP5
ECSIT
LRRC15
RNF168
CRYBG3
CISD2

RRBP1
UBXN2B
PRDX5
VDAC3
VDAC1
CASP7
TOM1
PRMT6
TMEM106A
CDC20
ACSL5
NDUFB6
TYSND1
NMT2
G0S2
CTSD
SLC4A7
RRP1B
SGMS1
SEMA3B
THOP1
SCARB1
ADSSL1
MRPS26
RBPMS2
ITPR2
AK2
BHLHB9
RFC4
OMA1
BCL2L13
CCDC30
SEMA4B
TMEM63B
NUP50
KAT2A
CSNK1A1
MFSD1
PUS7L
PLEKHF2
PEX6
UBAP1
DHRS3
DYNLRB1
PBK
MR1
EXOSC5
TNFRSF12A
ATF4
IFRD1
ARNTL
SIGMAR1
TSPAN5
CCL2
AMIGO1
ATAD3A
TXNIP
TRIP13
MAZ
DFNA5
ACADS
MED30
EPHX2
CORO6
ANXA5
KIF3B
CPT2
TOR1B
LSS
ZMYM1

BCL7C
TRPM4
ARHGEF40
UBE2D1
IDH3A
HSP90AA1
SEL1L
YWHAB
ENTPD5
ACSS2
FGFRL1
SUCLG1
EIF4B
RBKS
PHLDB1
SEC23A
MAGEH1
PRKCE
PGM2
RILPL2
ROCK1
TUBE1
PHF5A
RGS2
TAGLN
CREB3L2
RTN4
GFM2
NDUFB10
IVNS1ABP
PGP
NDUFV3
ACSS1
OPLAH
ST3GAL2
MRPL18
TUBA4A
FBXO21
CLSTN1
MYLIP
RPIA
PRDX3
LMAN2
CHCHD3
MFAP4
SLC11A2
H2AFJ
DHX32
DENND5A
TMEM150C
BDH1
COQ6
COTL1
CLDN15
SCYL2
DOCK9
UROD
DDX28
RBCK1
PPARA
ZDHHC8
PLXNA2
MGLL
FBXL15
DDIT4
NUS1
DNAJA2
NCS1
ATP5L
ATP1A2

NUDT9
BACE1
HERPUD1
BCL2L11
METTL9
ATP5O
TIMM9
UGDH
MAP2K5
LGMN
SLC5A6
SDPR
LAMB1
NUCB2
RARRES2
NOD1
ADAMTS1
JMJD6
PRKG1
DGKI
FAM136A
NFIX
AKAP1
HIBCH
SLC7A5
AIFM2
TIMM10
AK4
NLN
SORBS1
ETFDH
CYGB
NME7
UBE2C
NQO1
HSPA9
AXIN2
GSTK1
AP1G2
MAP3K2
SLC9A6
TSPO
HOXC9
RAB3D
TBPL1
ARRDC3
SDC1
MUL1
DDIT3
GGT7
HPCAL1
PSAT1
OSR1
GDPD3
PTGR1
FOS
MRPS5
CNTRL
CREG1
AGPAT2
HSPA5
TNFSF13B
OGFOD3
CMPK1
CYP27A1
PAFAH2
NPEPL1
H6PD
GNPNAT1
SAP30

SECISBP2L
HTATIP2
PEX3
PAMR1
PABPC4
HADH
FAM109B
ABCB10
ING4
IDI1
GLUL
TRAF4
HK2
ELOVL6
EPHX1
UBE2E1
DNAJC2
GADD45GIP1
SLC27A4
ASB6
PPIF
EHD3
COL14A1
TNFAIP8
EFEMP2
ABAT
CAV1
DGAT2
STRBP
HECTD3
TNFRSF1B
GTF3C4
LGALS9
SCARB2
IQSEC1
BAG2
HMGCL
TMEM164
SLC9A1
THBS2
ASNS
PPM1L
ISG20
KCNK15
HSP90B1
SMOC1
SPC25
SLC25A33
FAH
RCE1
UBE4B
TIMM8B
STAP2
TRIM44
DNTTIP1
TRAF1
GRN
FAM98C
SLC2A4
TRIM32
LDLRAD3
LRRC59
ADCY6
NPR2
TMEM170B
AP3S1
STOML2
INMT
LAS1L
FLAD1

LNX2
ERP44
PKN1
ITM2A
RHBDF2
CEBPA
ACSF3
JMJD1C
TSC22D4
OLFML2B
S100A10
WDR76
MRPS2
CDKN1C
SIRT3
ST6GALNAC6
THUMPD2
ACTB
TRMT12
TLR4
CMBL
RAB30
OSBPL11
PCDH9
GBF1
ACVR2B
ZSWIM7
SERPINE1
UBQLN1
SOCS2
GBP3
PPP2R2D
TIMM44
ILVBL
OXNAD1
SMIM20
PLEKHH3
PRELID1
FAM162A
INSR
TGFBR2
INTS4
GALE
NT5C2
CIZ1
WWTR1
CIB1
OXSR1
TNFAIP2
RAD51D
MTSS1
ADPRM
RGP1
DLD
CDK5
MMD
CUL2
DENND4C
BRPF3
PKM
MORC4
ETV5
LY6E
SCUBE2
GSTO1
NRCAM
LSM1
LYRM4
IDUA
CSRP2

HIPK2
CCND2
ADIPOR2
PEX11A
CDC34
PKDCC
ECHDC2
UPP1
CISD1
TMX2
PPM1F
PCSK5
NDUFA9
GMNN
SUDS3
RNF149
EGLN3
PEX19
UNC119
GPX3
NOV
PSMD7
SSX2IP
RAD21
LRP4
TMTC4
SNRPB
TAPT1
ENC1
MICALL2
POP5
TCEA3
LPCAT3
SF3B4
VOPP1
GTF2IRD1
SMARCA2
ACOT4
STC2
COL6A1
FOXD1
GPRC5B
F11R
ACTA2
DERL1
F3
MAP4K4
PLEKHM2
SYNPO
DLAT
TRIT1
LMNA
HOXC6
PRICKLE1
PEG10
CCDC92
KLF10
FOXO3
PARD3B
LPAR1
PPA1
CEBPB
HMGA1
POPDC3
GEM
TFPI2
NANOS1
PDHB
SHKBP1
JUN

NAGK
KDM3A
ATP2A2
RETSAT
ADAM15
FAM110B
CALHM2
FBXO32
PNN
TGFBI
CLPP
PPP1R3G
TRIM25
NUP62
RASA3
HNRNPL
PGD
ITPRIP
E2F3
ABHD6
CUTA
WAC
ATPAF2
PTPN13
ECD
FDPS
SLC52A2
SUCLA2
MTMR11
SUOX
IDH3B
NDUFS7
SELENBP1
FOSL1
NPC1
PRKCA
ALDH9A1
STK17B
B3GAT3
ATP8B1
MAT2A
LRP11
PGPEP1
MBOAT2
SEMA3C
DPYSL3
S100A13
PAK2
PRKAG1
IFI44
SMAD5
GARS
FAM49B
PNPLA2
BCKDHB
PNKD
HDAC6
UBC
PSMB9
CBR1
ZNHIT6
PIK3IP1
RXRB
ADIPOQ

**Supplementary Table S2: PPRE-Driven Genes from Lemay and Hwang Manuscript, Identified in Podocyte and Adipocyte Datasets**

| Podocyte Not Adipocyte (139) | Podocyte And Adipocyte (478) | Adipocyte Not Podocyte (22) |
|---|---|---|
| ZNF691 | ZNHIT2 | ZIC1 |
| ZNF485 | ZNF668 | TIMP4 |
| ZMAT1 | ZNF608 | STMN2 |
| ZIC2 | ZNF462 | RPS17 |
| ZBTB9 | ZNF436 | RNF5 |
| ZBTB12 | ZNF416 | PRRX1 |
| XK | ZNF219 | PDGFRA |
| WNT3 | ZMAT5 | LSP1 |
| WNT10A | ZMAT2 | LRRC4 |
| WIBG | ZHX3 | LMO3 |
| WFIKKN1 | ZFHX4 | LINS1 |
| UCP3 | XRCC5 | INHBB |
| UCKL1 | WTAP | GPC3 |
| TRIM3 | WDR46 | FYB |
| TRERF1 | WDR27 | FGR |
| TESK2 | WDR18 | EYA1 |
| TENC1 | VPS41 | CXCR4 |
| SYT7 | VPS29 | COL1A2 |
| SYT12 | VPS16 | CD1D |
| SYN1 | VLDLR | CCR1 |
| STAG3 | VEGFB | AQP7 |
| STAC3 | VAT1 | ACE |
| SPTBN2 | USP37 | |
| SPOCK3 | UMPS | |
| SPIB | UCK2 | |
| SPAG8 | UBE2E1 | |
| SP6 | UBE2C | |
| SOX6 | TXNIP | |
| SLC7A14 | TUB | |
| SLC39A5 | TTLL5 | |
| SLC25A34 | TSPYL2 | |
| SLC23A3 | TSEN34 | |
| SLC14A1 | TRPS1 | |
| SGK2 | TRPM4 | |
| SERTAD3 | TRIM65 | |
| SEPT6 | TRAPPC3 | |
| SEMA4G | TPP1 | |
| SEMA3E | TP53INP1 | |
| SCN4B | TP53BP1 | |
| RASGRF2 | TOP2B | |
| RAB24 | TOM1L2 | |
| PTP4A3 | TOB1 | |
| PSMA6 | TNRC6A | |
| PSD | TNPO3 | |
| POU5F1 | TNK2 | |
| POLD4 | TNIP1 | |
| PITX3 | TNFRSF1B | |
| PITX2 | TMEM53 | |
| PHYHIP | TMEM43 | |
| PFKFB1 | TMEM30A | |
| PEX5L | TLR4 | |
| PDZK1 | TLK1 | |
| PDF | TIMELESS | |
| PCDH17 | THAP8 | |
| PAX6 | TGFBR2 | |
| PAX2 | TERF2IP | |
| OSCAR | TEAD3 | |
| OBSCN | TCTA | |
| NTN1 | TBX15 | |
| NRK | TBL1XR1 | |
| NR1I3 | TAX1BP1 | |
| NETO1 | TARBP2 | |
| MYL1 | TAPBP | |
| LRRC29 | TAOK3 | |
| LMX1B | SYNGR2 | |
| LLGL2 | SYNGAP1 | |

| | |
|---|---|
| LIPC | SYNE1 |
| LHX2 | STOML2 |
| LAG3 | STMN1 |
| KIRREL3 | STK19 |
| KIF5A | STK11 |
| KCP | STEAP2 |
| KCNN4 | STAT5A |
| KCND1 | SSR4 |
| ITGB1BP2 | SRR |
| ISL2 | SRP72 |
| IRF2BP1 | SRC |
| IQCD | SP2 |
| INCA1 | SOCS2 |
| IL4I1 | SNRPD2 |
| HUNK | SMYD5 |
| HSPB8 | SMTN |
| HSD17B7 | SMARCD2 |
| HPX | SMARCD1 |
| HOXD3 | SMARCA1 |
| HOXD1 | SMAD3 |
| HOXC4 | SLC6A9 |
| HOXB9 | SLC4A2 |
| HOXA3 | SLC41A1 |
| HOXA13 | SLC36A4 |
| HIST1H2AI | SLC35E1 |
| GUCY1B2 | SLC35A4 |
| GPT | SLC31A2 |
| GPR64 | SLC2A4RG |
| GPR124 | SLC27A3 |
| GPR115 | SLC26A6 |
| GLI1 | SLC25A28 |
| GBX2 | SLC25A25 |
| GAS2 | SLC25A13 |
| FUT1 | SLC22A17 |
| FTSJ2 | SLC1A4 |
| FGFR4 | SLC1A3 |
| FBXO6 | SKIV2L |
| FANCB | SIRT7 |
| EPN3 | SIRT6 |
| EEF1G | SIPA1 |
| DSG3 | SFXN5 |
| DOK3 | SEZ6L2 |
| DND1 | SERP1 |
| DNAH3 | SENP3 |
| DLG2 | SEMA7A |
| CYP27B1 | SEMA3A |
| CYP26B1 | SEC24C |
| CYP1A1 | SDSL |
| CXXC4 | SDHD |
| CXCL16 | SDHC |
| CPT1B | SCRIB |
| CNIH2 | SCNM1 |
| CLDN3 | SCML1 |
| CIDEB | SCAMP3 |
| CHRFAM7A | SAT2 |
| CHMP4C | SAMD1 |
| CELSR3 | RTN3 |
| CAPN3 | RTN2 |
| CACNG6 | RREB1 |
| C9orf37 | RRAS |
| C1QL4 | RPS6KA2 |
| C10orf55 | RPS26 |
| BSCL2 | RPS2 |
| BDNF | RPL9 |
| BCKDHA | RPE |
| BAI2 | RORA |
| ARHGAP30 | RNPEP |
| AQP3 | RNF182 |
| AMIGO3 | RNF14 |
| AFF3 | RIT1 |
| ADAM8 | RILP |
| ABCB6 | RHOBTB1 |

| | |
|---|---|
| A1BG | RFFL |
| | REST |
| | RCOR3 |
| | RCL1 |
| | RBM8A |
| | RBM15B |
| | RBKS |
| | RASAL2 |
| | RARA |
| | RAD23B |
| | RAD21 |
| | RAD17 |
| | RAD1 |
| | RAB5C |
| | RAB11FIP2 |
| | QARS |
| | PTX3 |
| | PTRF |
| | PTPN23 |
| | PTPN14 |
| | PTMS |
| | PTEN |
| | PSME2 |
| | PSME1 |
| | PSMC5 |
| | PRR3 |
| | PRKCD |
| | PRKAG1 |
| | PRKACA |
| | PRKAA2 |
| | PRDM1 |
| | PRAF2 |
| | PPP2CA |
| | PPP1R12B |
| | PPP1R10 |
| | PPOX |
| | PPARGC1A |
| | PPARG |
| | PPARA |
| | POLR2H |
| | POLR2A |
| | POLR1D |
| | POLK |
| | POLG2 |
| | POFUT1 |
| | PLEKHF2 |
| | PLEKHA4 |
| | PLCD3 |
| | PLCB3 |
| | PKP4 |
| | PITPNM1 |
| | PIK3C2B |
| | PICALM |
| | PHTF1 |
| | PHLDB3 |
| | PHLDB1 |
| | PHF5A |
| | PGGT1B |
| | PFN1 |
| | PFDN2 |
| | PDLIM2 |
| | PCOLCE |
| | PCGF2 |
| | PCDH7 |
| | PCBP4 |
| | PARVA |
| | PAQR4 |
| | PANK2 |
| | PAM |
| | PAK1IP1 |
| | PAK1 |
| | OSGEP |

NXF1
NUP188
NUMBL
NUDT8
NUDT6
NUDCD1
NR4A3
NR4A1
NR2F6
NR2F2
NR2F1
NR2C2
NR2C1
NPHP4
NPAS1
NOVA1
NMNAT1
NISCH
NFIA
NEU1
NEDD9
NDUFB11
NCOA2
NBEA
MYO1C
MYL6
MYH10
MYADM
MVP
MTMR4
MST1
MRPS18C
MRPL9
MRPL48
MRPL24
MRPL21
MRPL14
MOV10
MMP19
MMP15
MLX
MLF2
MITF
MID2
MEF2D
MEF2C
MED8
MDM2
MDM1
MCEE
MCAM
MARVELD1
MAPRE2
MAPK8IP1
MAPK11
MAML1
MADD
LZTR1
LTBP2
LRP1
LRFN3
LPL
LNX2
LNPEP
LMCD1
LIX1L
LASP1
KPNA4
KLHL18
KLF4
KLF12
KLF11

KIRREL
KIAA1715
KIAA0319L
KHSRP
KCTD9
KCTD12
KCTD11
KCNK6
KCND2
JUNB
ITGA3
IRAK1
IQCG
INPP5B
ING4
ING1
IL11RA
IGFBP6
IFITM2
ICMT
HSPBP1
HPCAL1
HOXD8
HOXB4
HOXA6
HOXA10
HMGA1
HINT2
HIC2
HIC1
HEBP2
HDAC3
HDAC1
HADHB
H2AFY2
H2AFX
GTF2A1
GSN
GRWD1
GRK4
GPS2
GPRC5C
GNB1L
GNAS
GNA13
GMPPA
GLCE
GIT2
GIT1
GFRA1
GDPD3
GCLM
GCAT
GBA2
GALT
GALNT7
GADD45A
GABPA
GAB1
G0S2
FZD2
FURIN
FOXP1
FOSL1
FLOT1
FLNC
FLI1
FKBP5
FHL3
FBXL17
FBXL12
FBN1

FASTK
FANCD2
FAM50A
FAM35A
EWSR1
ETV2
ETFA
ERMAP
EPN2
ENTPD5
ELOVL3
EIF4G1
EIF2B4
EGLN2
EEF1D
DVL2
DUSP6
DST
DPM2
DPF3
DNAJC17
DNAJC13
DNAJA2
DMPK
DMD
DLG4
DLEU1
DHRS1
DDIT3
CUEDC2
CSNK2A2
CSAD
CRELD1
CREB5
CREB3
CRABP2
CPNE2
CPEB2
CORO1B
COPG2
COL1A1
CNOT2
CNOT1
CLN5
CLDN15
CLCF1
CIC
CHPF
CHML
CHEK1
CHCHD4
CEBPB
CDKL5
CDK5
CDIPT
CDCA5
CDC6
CDC42EP3
CD9
CD68
CD3EAP
CD36
CD2BP2
CCT7
CCND1
CCHCR1
CCDC18
CBX5
CAMK2N1
CACNB2
C6orf136
C5orf15

C21orf58
C2
C1orf43
C11orf31
BUB3
BRPF3
BRF1
BRE
BMF
BLOC1S2
BCORL1
BCL9L
BCL7C
BAZ1B
BANK1
BAHD1
BACH1
ATPAF1
ATP8B2
ATP6V1F
ATP6V1A
ATP6AP1
ATP1B1
ATP10A
ATN1
ATAD3B
ASB6
ARPC5
ARIH2
ARID3A
ARID1A
ARHGAP5
ARHGAP22
ARCN1
AP1G2
ANKRD12
ANAPC11
ALDOC
ALDOA
AKT1S1
AKAP8L
AGPS
AGL
ADCK4
ADAM12
ACSL3
ACRC
ACP2
ACADVL
ACAA2
ABL1
ABCF3

### Supplementary Table S3: PPRE-Driven Genes in the PPARGene Dataset and Lemay and Hwang Manuscript

| PPARGene Dataset | Lemay & Hwang | Both |
|---|---|---|
| AACS | DPM2 | PAX7 |
| ELMO2 | RPS17 | HMGCS2 |
| KRTCAP2 | ZHX3 | HIC2 |
| ABRA | BBOX1 | PDLIM2 |
| PYROXD2 | PDF | PHYHIP |
| FAM187A | ZCCHC16 | GRM2 |
| BTG2 | CXCR4 | RCL1 |
| DNAJB8 | IRAK1 | SYNGAP1 |
| NDUFS6 | ADAM12 | NMNAT1 |
| XDH | POLK | ACSL3 |
| 1700024G13 | PAK1 | ALDOC |
| TACC2 | MITF | RAD23B |
| PDHA1 | ZBTB12 | ITGB1BP2 |
| SLC4A1 | FLJ10385 | LLGL2 |
| ABCB7 | CD2BP2 | GCLM |
| HSPB1 | FGR | PCDH7 |
| PFKP | EPN2 | PTRF |
| PBX4 | KCNK6 | MMP15 |
| ARRB2 | XK | SRP72 |
| DECR1 | RNF14 | CYP1A1 |
| CBR2 | BANK1 | SLC39A5 |
| PRADC1 | SLC13A1 | MYH2 |
| CMTM8 | B3GAT1 | STAT5A |
| CRTAM | STGC3 | KLF4 |
| CSPG4 | HOXA13 | NR1I3 |
| ACOT12 | PCOLCE | PPARGC1A |
| IGF1 | NFIA | ETFA |
| SLC35G3 | C5orf19 | PFKFB1 |
| ITPK1 | BLOC1S2 | SCN4B |
| G3BP1 | TAOK3 | PPOX |
| HSD17B10 | ITGA3 | ACADVL |
| LMAN1 | POU3F1 | GLI1 |
| TMEM139 | OR8D4 | SDHD |
| CYP4F16 | USP52 | SLC25A34 |
| UQCR11 | PHLDB3 | H2AFY2 |
| GDF15 | BAZ1B | INCA1 |
| CYCS | AGBL4 | SGK2 |
| WISP2 | ZIC2 | FKBP5 |
| GPSM2 | LOC492311 | FLI1 |
| CPXM2 | TERF2IP | PTP4A3 |
| TBX10 | FLJ23518 | HSD17B7 |
| NOS2 | COL1A1 | PAK1IP1 |
| GYS1 | ZFHX1B | VAT1 |
| TRIB3 | PRG | SENP3 |
| TAF9B | C1orf61 | ACAA2 |
| DTNBP1 | CLN5 | TMEM53 |
| REEP6 | UNQ2446 | MRPL9 |
| CLUH | GNAS | SLC23A3 |
| ZG16 | PYY | GPR81 |
| PROP1 | KCNC1 | GNA13 |
| CDK14 | SIRT6 | HUNK |
| NDUFS1 | CCL19 | GPT |
| SOD2 | FLJ21616 | SLC26A6 |
| IGBP1B | KIAA0339 | UCKL1 |
| AES | SLC27A3 | BMF |

| | | |
|---|---|---|
| ARL2 | KIAA1715 | ZIC5 |
| SYCP3 | FGF17 | SDSL |
| KCNK3 | EIF3S2 | LRP1 |
| NDUFAF4 | CABP1 | PSD |
| STK16 | PLIN | CLDN3 |
| GDA | NR2E3 | NR4A1 |
| AQP9 | PRDM16 | AQP3 |
| NDUFAB1 | RBM21 | IGFBP6 |
| SLC15A2 | SIPA1 | MRPL14 |
| FGFR1 | CTSL2 | AGL |
| MREG | CYorf15B | TBL1XR1 |
| AGPAT1 | CDIPT | BCKDHA |
| CERK | SYT12 | EGLN2 |
| ATP5D | PCBP4 | HADHB |
| UNC5A | DKFZp761A | BAHD1 |
| RAB40B | MAPBPIP | TNIP1 |
| SERPINA12 | GMPPA | POLR2A |
| MRPL38 | LOC402682 | FGF10 |
| PIR | MGC17299 | NR2F6 |
| LRRN1 | DEFB119 | PCDH17 |
| LONP1 | FLJ40201 | SMTN |
| PPCS | GUCY2C | GALT |
| MRPL1 | FLJ46020 | NR2C1 |
| AMY1 | CNOT1 | SLC41A1 |
| ARL6IP1 | ZNHIT2 | LPL |
| BET1L | ADCK4 | ADRB3 |
| PTCD3 | FYB | XRCC5 |
| FIBCD1 | CDC42EP3 | POU5F1 |
| FADS1 | PTMS | FASTK |
| TMEM184A | FLJ23129 | PSMA6 |
| TGOLN1 | ZMAT5 | UCP3 |
| PCYT2 | HTR3A | PSME2 |
| SERPINF1 | C20orf118 | TBX6 |
| TSC22D1 | FGF6 | MMP19 |
| ATXN10 | CNTFR | RHOBTB1 |
| ECH1 | ATAD3B | VEGFB |
| RHOT1 | WNT3 | UCK2 |
| FAHD1 | LGI4 | MEF2D |
| 4833439L19F | KIBRA | NFE2 |
| ELOVL5 | PB1 | KLF11 |
| EPS8 | CHRM1 | NR2F1 |
| A130010J15F | CDC6 | ARHGAP30 |
| DIP2A | NYD | CREB3L3 |
| OXT | CLDN6 | SMAD3 |
| SHC1 | CCDC18 | ELOVL3 |
| CCL22 | PICALM | MYO1C |
| ACTR8 | AGPS | BDNF |
| HSD11B2 | NEU1 | SEMA3E |
| BRCA1 | NAG6 | CHEK1 |
| GEN1 | NR2C2 | CD36 |
| LRFN5 | C9orf25 | KCNH2 |
| FAM96B | MGC11102 | PPARG |
| S1PR2 | GPRC5C | ISL2 |
| PAPSS2 | TENC1 | DMPK |
| A2M | OSGEP | SMYD5 |
| PIK3R5 | ETV2 | MYL6 |
| HIGD1B | ATP10A | IFITM2 |
| USP6NL | PRRX1 | LTBP2 |

| | | |
|---|---|---|
| GRPEL1 | PAQR4 | EDN3 |
| SLCO2B1 | HDAC3 | VLDLR |
| KRT1 | C6orf27 | NUDT6 |
| ROCK2 | FLJ45831 | ATN1 |
| PPP1R15B | HAPLN4 | SLC4A2 |
| ZCCHC5 | TIAF1 | G0S2 |
| ZMYND15 | CATSPER4 | TRERF1 |
| HES1 | HIG2 | RNF5 |
| A1CF | RNF182 | PLEKHF2 |
| CAR6 | SCAMP3 | TXNIP |
| CLDN16 | CELSR3 | BCL7C |
| ISL1 | GRWD1 | TRPM4 |
| C1QTNF3 | UMPS | ENTPD5 |
| RIC8 | RAD1 | RBKS |
| VGLL3 | ERMAP | PHLDB1 |
| SEL1L3 | LOC441046 | PHF5A |
| PDCD4 | WNT6 | C1QTNF4 |
| CEP350 | TSP50 | CBLN1 |
| HSPA1L | TRIP | OSCAR |
| FAM73A | TLK1 | CLDN15 |
| RHOC | LOC112476 | PPARA |
| PPP3CA | KIAA0404 | PCDH1 |
| DDB2 | WFIKKN1 | DNAJA2 |
| NEIL1 | SERP1 | EPN3 |
| TSPAN13 | EPB49 | FABP1 |
| FITM2 | CREB3 | PACSIN1 |
| SPIRE2 | CA9 | UBE2C |
| TRF | GPR174 | AP1G2 |
| MKNK1 | TCTA | GRIK5 |
| STARD5 | KIAA0319L | DDIT3 |
| DNAJC15 | KIAA0377 | HPCAL1 |
| CCL9 | SAT2 | GDPD3 |
| MGAT4B | KRTAP13 | ING4 |
| NCOR2 | BHLHB2 | UBE2E1 |
| NEPN | ATP6V1A | ASB6 |
| TRP53INP2 | INPP5B | TNFRSF1B |
| IL17RC | DND1 | NEUROG2 |
| LRRC8D | LOC541469 | STOML2 |
| GSTA2 | C1orf66 | LNX2 |
| ALDH18A1 | ATBF1 | CYP26B1 |
| CEP170 | VPS16 | TLR4 |
| UFM1 | CCND1 | SOCS2 |
| ACAA1A | CA5A | TGFBR2 |
| TPI1 | IL4I1 | CDK5 |
| CORO2A | DSCR2 | PADI6 |
| MAOA | RRAS | BRPF3 |
| APOE | MGC13053 | PDZK1 |
| ANGPT4 | SYT7 | RAD21 |
| VAT1L | PGGT1B | APOA1 |
| ALDH2 | CSNK2A2 | KLK11 |
| GMPR | PTPN14 | CEBPB |
| IP6K1 | MGC14288 | HMGA1 |
| EMP2 | PRDM13 | FOSL1 |
| FIGF | ZNF536 | PRKAG1 |
| SRSF6 | ZNF385 | AQP7 |
| QDPR | AMIGO3 | |
| MRAP | FGFR4 | |
| HIST2H2AA1 | PPP1R14D | |

| | |
|---|---|
| HSPD1 | OR2L13 |
| EIF1 | SLC31A2 |
| ZFP219 | GRIN2A |
| RIN2 | PAX2 |
| TMEM98 | MGC26694 |
| CDK1 | NR4A3 |
| SLC38A10 | BCL11B |
| FRRS1 | LMX1A |
| APOC4 | GLCE |
| OPHN1 | GPR124 |
| NDUFA8 | MRPS18C |
| SHBG | GBX2 |
| FABP5 | C15orf17 |
| SLC16A5 | FLJ10099 |
| GCC2 | PCYT1B |
| GLTSCR2 | BLK |
| TOR3A | CNOT2 |
| PLIN2 | ARL6IP |
| CHMP1B | ABI3 |
| CD82 | IPLA2 |
| PPM1K | ZNF663 |
| PMM1 | POLD4 |
| CPEB3 | WNT16 |
| NFKB2 | LOC55565 |
| BCL2 | CDCA5 |
| FZD9 | STMN1 |
| ADI1 | LIX1L |
| SCP2 | C20orf28 |
| KL | LOC220686 |
| ALCAM | CHES1 |
| SH2B2 | HTR3D |
| TRIM29 | CHMP4C |
| ID1 | CD68 |
| OBP2A | SMARCA1 |
| COL18A1 | LOC147650 |
| ATP1B4 | SSR4 |
| MPP4 | NR2F2 |
| CDS1 | RPS6KA2 |
| ERLIN2 | PRDM1 |
| PPP1R9A | FLJ36874 |
| TIMM22 | C20orf112 |
| PDZRN3 | P2RY4 |
| GPD1 | WIBG |
| ADAM19 | ZNF608 |
| ADAM32 | GBA2 |
| CES1F | CCT7 |
| PRPS2 | FLJ22457 |
| GCK | VPS29 |
| MYBPC1 | RAB24 |
| ADRA2A | FLJ10276 |
| CELA1 | PTPN23 |
| ALAD | C4B |
| ASS1 | ATP6V1F |
| GSTM4 | FLJ11016 |
| F7 | BUB3 |
| RGS4 | VIL1 |
| UQCRC1 | LECT1 |
| SNX6 | ASB4 |

| | |
|---|---|
| PPP2R5A | PAM |
| SAMM50 | C10orf55 |
| MRPL34 | CCL23 |
| 4931406C07I | HOXD8 |
| ESRRA | FLJ25037 |
| FNDC8 | KPNA4 |
| LDHB | ZNF278 |
| LRRC8A | SIRT7 |
| GNA11 | ALOXE3 |
| DDX49 | RPE |
| TLCD2 | C1orf22 |
| EBPL | HTR4 |
| RHBDF1 | EIF2B4 |
| SNX25 | CRABP2 |
| IGSF11 | LOC201725 |
| RASIP1 | GATA3 |
| KCNE3 | FLJ43860 |
| CPEB1 | ENDOGL1 |
| ERO1L | PIK3C2B |
| ACOT7 | PLA2G12B |
| H2-Q10 | PDZK2 |
| ENDOG | KLHL1 |
| EZR | RTN2 |
| SLC38A4 | GADD45A |
| ABHD12 | HIC1 |
| HES7 | PX19 |
| LRRTM3 | GP1BB |
| PTGES | TNRC6A |
| PEX16 | CYP27B1 |
| FGF1 | ZNF691 |
| WDR6 | PAX6 |
| FUT4 | C10orf9 |
| CD300A | KIAA0882 |
| XRCC6BP1 | LOC152485 |
| DNAJA3 | MDM2 |
| ITFG2 | NETO1 |
| EPHA4 | HOXB4 |
| TMEM11 | ICMT |
| RARB | NBEA |
| IMMT | LINS1 |
| KCNQ5 | MAML1 |
| UBE2L6 | ZMAT1 |
| UGT2B36 | MYL3 |
| AGGF1 | FLJ36208 |
| EPGN | EEF1D |
| NPHS1 | CHML |
| TPBG | MTMR4 |
| SPRY2 | NUDT8 |
| LZTFL1 | GPR144 |
| TRPV5 | C16orf47 |
| GSTP1 | HOXC4 |
| GMFB | TBC1D21 |
| LTBR | MGC2744 |
| AGTPBP1 | GUCY1B2 |
| PRR5 | NISCH |
| TLE2 | MGC5509 |
| PCSK6 | ISGF3G |
| RPRD1B | TMEM43 |

| | |
|---|---|
| 1110034G24 | FLJ22573 |
| 2310079G19 | COPG2 |
| FKBP4 | PH |
| ADRA1A | FLJ33817 |
| PNPLA6 | SLC35E1 |
| GIPR | KCTD11 |
| ZFP385A | NELF |
| KISS1R | CKMT1A |
| NRG4 | ABL1 |
| FAM154B | PME |
| PERP | DKFZp434H2215 |
| SCN11A | KLHL18 |
| YIPF2 | SP2 |
| FHL1 | KIAA1967 |
| CDH24 | CHRFAM7A |
| TNFSF12 | TCF1 |
| PBX1 | TMP21 |
| CSRP3 | TARBP2 |
| PSMD5 | SYNE1 |
| LACTB2 | LOC440503 |
| FAM126B | C2 |
| LCA5L | E2IG5 |
| RBPMS | MGC13102 |
| GANC | CRELD1 |
| CHRNA2 | SNCG |
| ACO2 | MGC21518 |
| TPMT | RNPEP |
| ELFN1 | MGC21830 |
| KCNA4 | C14orf68 |
| LOXL1 | WDR27 |
| ARID5A | FAM72A |
| NFE2L2 | FBXL17 |
| EML5 | SPTBN2 |
| FAM195A | RPL9 |
| IL23R | OTOP3 |
| SH2D2A | CPLX2 |
| SETD8 | ARHGAP5 |
| ACAD11 | GAS2 |
| OLFR555 | CACNB2 |
| ACSF2 | FLJ25421 |
| HP | SRC |
| TJP3 | FLJ23375 |
| PFKL | TIMELESS |
| SNN | GRK4 |
| TMEM140 | PPP2CA |
| GABBR1 | BRE |
| CHRM4 | PHF15 |
| TMEM25 | MAPRE2 |
| NIPAL4 | SOX6 |
| PROX1 | MGC13045 |
| POLD2 | MAPK11 |
| CAPN6 | PHTF1 |
| LSM3 | N/A |
| PPL | DKFZP761H1710 |
| ACOT2 | HOXD3 |
| RRAD | AKAP8L |
| AHRR | NDUFB11 |
| SNAI1 | STK19 |

| | |
|---|---|
| RASSF4 | PANK2 |
| MTHFD1 | SLC1A3 |
| SOX9 | BF |
| GJB5 | ADAM8 |
| CASC4 | PDZK7 |
| CRABP1 | HLXB9 |
| 6330416G13 | LRRN5 |
| SLC35F1 | STK11 |
| SYTL4 | SPAG8 |
| PRR36 | NOD27 |
| CEBPD | PSMC5 |
| ADRB1 | EGFL4 |
| VRK2 | C6orf49 |
| SNTG2 | SLC2A4RG |
| JUND | DNAH3 |
| MOSPD1 | CXorf1 |
| PHB2 | eIF2A |
| EHBP1L1 | LIPC |
| NDRG2 | KIRREL3 |
| CAMK2B | GABRG2 |
| CELF5 | C9orf100 |
| LIN28B | NCOA2 |
| ZFP568 | DNAJC17 |
| COG4 | MGC2803 |
| NDUFB8 | KIRREL |
| DNAJC11 | GPC3 |
| DIXDC1 | PDGFRA |
| TIMM17A | CACNG1 |
| CADPS2 | MGC21644 |
| SEMA3G | FLJ40448 |
| NGFR | DMD |
| EEPD1 | ZNF668 |
| GRHL1 | ABCB6 |
| KLHL10 | ZMAT2 |
| OLFR231 | PRKACA |
| CCDC132 | NOVA1 |
| AMMECR1 | SEC13L1 |
| LRRC2 | MLL2 |
| DLST | HDAC1 |
| NEDD4L | ANKRD33 |
| ANO9 | LMCD1 |
| NCAPD3 | MYCL1 |
| IL1A | GFRA1 |
| FST | ZNF161 |
| UPK3B | KCND2 |
| TMEM79 | CDKL5 |
| AQP8 | SLCO5A1 |
| LDLR | RFFL |
| SCRN1 | RBM15B |
| PROSER2 | FCER1G |
| PMVK | FLJ36031 |
| COQ5 | TOP2B |
| SGK1 | TESK2 |
| DDI2 | DHRS1 |
| SIX2 | HOXA6 |
| RNF7 | TRAPPC3 |
| POLR3G | HAMP |
| CPD | PLVAP |

| | |
|---|---|
| DCT | GRK7 |
| FBF1 | FLJ37794 |
| IRF7 | ZNF297 |
| PELI3 | TIMP4 |
| TBC1D8 | FBXO6 |
| HSPA2 | CACNG6 |
| SLC38A9 | C3F |
| KRT8 | FLJ10925 |
| ACAP3 | PTCHD1 |
| NFKBIZ | CIDEB |
| RAB20 | ARCN1 |
| CAR2 | MR |
| GATSL2 | TTLL5 |
| TAF9 | STMN2 |
| AGTR1A | NRK |
| DGAT1 | 6-Sep |
| METTL5 | SFXN5 |
| ISCU | RP11 |
| MYCL | CHPF |
| RAB15 | MID2 |
| K230010J24R | COL1A2 |
| PHGDH | GPS2 |
| HS3ST6 | GRCC10 |
| MIP | FY |
| FDX1 | FAM13A1 |
| F2 | KIAA0467 |
| CASP14 | VMD2 |
| XIAP | ARID1A |
| SH3BGRL2 | MGC35366 |
| WNT5A | DKFZp564I122 |
| APOA2 | KCP |
| TRABD2B | C1orf43 |
| DOPEY2 | CD3EAP |
| PEX26 | MAPK8IP1 |
| PTGS2 | FANCD2 |
| SLC29A3 | EFS |
| LGALS4 | PITPNM1 |
| RAB11FIP4 | MYADM |
| IMPAD1 | LZTR2 |
| SQLE | MEF2C |
| ADAMTSL4 | FAM50A |
| OPN3 | AFF3 |
| OLR1 | LOC340529 |
| DAGLB | ZNF416 |
| HCFC1R1 | ZNF485 |
| NRAP | SPIB |
| GLIPR2 | GNG8 |
| SLC7A4 | C6orf114 |
| WBP11 | MGC42718 |
| TRIM63 | QARS |
| TEX2 | KIAA0553 |
| LYPD3 | HSPB8 |
| MAP1A | FBXL12 |
| MAP4K3 | PTEN |
| 2610318N02 | MGC11271 |
| COX7B | RLBP1 |
| TMEM185B | GTF2A1 |
| ANKRD17 | FLJ10404 |

| | |
|---|---|
| SLMO2 | LZTR1 |
| GRPEL2 | ATP1A3 |
| SNTA1 | BTLA |
| IL21R | CAPN3 |
| CACNG2 | FLJ22875 |
| TLR5 | LOC120379 |
| FAM78A | FHL3 |
| NT5E | POLG2 |
| FOXN2 | ASCL3 |
| CST11 | CD1D |
| CD200 | SMARCD1 |
| GALNT9 | PAX4 |
| FAM129B | ZNF219 |
| GCGR | TETRAN |
| ITGA6 | BTBD5 |
| NCK2 | POU3F4 |
| SLC16A2 | SP6 |
| SC5D | THY28 |
| TRIM35 | KCND1 |
| USP11 | LNPEP |
| EIF4E1B | TRDN |
| TMEM72 | MGC29814 |
| PRKDC | ANKRD15 |
| IMPA2 | TSGA14 |
| SULF2 | ATP8B2 |
| KLRG1 | DVL2 |
| NEURL3 | APOC3 |
| S100B | NPAS1 |
| NET1 | TRPV6 |
| LAMC3 | UNQ501 |
| TOX | ACRC |
| MVD | MRPL24 |
| MGP | ATP1B1 |
| CHDH | C5orf15 |
| CLDN10 | C9orf37 |
| ACLY | SEMA3A |
| DNAJC1 | TRIM3 |
| ARHGAP12 | C14orf121 |
| LGALS12 | REST |
| GABRG3 | RTN3 |
| MEX3D | 1-Nov |
| CCNDBP1 | MTP |
| SCCPDH | FZD2 |
| GPX4 | PARVA |
| CCDC85B | TEAD3 |
| PRMT1 | ARID3A |
| CILP2 | ALS2CR2 |
| ITIH3 | TLX2 |
| TMOD2 | FLJ40137 |
| MAB21L2 | ZNF286 |
| PPDPF | SFRS16 |
| FAM107B | MRPL48 |
| IRS2 | FLJ12787 |
| ALG9 | SYN1 |
| PIM3 | CCHCR1 |
| MEGF11 | MED8 |
| ANKRD40 | LRFN3 |
| COMTD1 | PLCB3 |

| | |
|---|---|
| ACOT9 | IMP |
| GPX2 | RAB11FIP2 |
| ROPN1L | LRRC4 |
| SPATS2L | RCOR3 |
| ADCK2 | ACP2 |
| SLC48A1 | ACPT |
| ERC2 | LAG3 |
| RBP7 | PRAF2 |
| KLK12 | FLNC |
| PDE3A | THAP8 |
| UQCRQ | CUGBP2 |
| RIMS1 | CDX2 |
| RCN1 | C4A |
| KAT5 | GRIN2C |
| BCAR3 | SLC7A14 |
| GAL3ST1 | PRKCD |
| MAP3K6 | CALCA |
| DHCR7 | VPS41 |
| TPD52L1 | TRIM65 |
| SPATC1 | MGC24133 |
| SCD1 | GNB1L |
| CRAT | BCL9L |
| HN1 | TAPBP |
| POR | GPR64 |
| VEGFA | RP42 |
| TRUB1 | KRTAP15 |
| CSRNP1 | C1orf51 |
| TRIM14 | HOXA3 |
| ELL | TOM1L2 |
| FAM89A | PHOX2B |
| CYP4A14 | CPNE2 |
| D630003M21 | NR2E1 |
| AIF1L | FLJ10159 |
| AREG | ARIH2 |
| NRSN1 | SRR |
| FATP1 | CORO1B |
| CYP2A4 | BRF1 |
| 1700011L22R | ARPC5 |
| SHANK3 | SYNGR2 |
| ACKR2 | RPS2 |
| EEF2K | SEC14L3 |
| TUFM | EYA1 |
| MPDU1 | CUEDC2 |
| MYC | PITX2 |
| ABHD16B | FLJ45187 |
| FZD10 | MGC11134 |
| DECR2 | ZFHX4 |
| GYG | ADAMTS19 |
| SPTLC3 | TSPYL2 |
| MUS81 | SLC25A13 |
| PHYH | LRRC29 |
| DAG1 | MIG |
| TIMP1 | MARVELD1 |
| RUVBL2 | DLG4 |
| CYP2B10 | ZIC1 |
| PFN4 | SPOCK3 |
| 42430 | FURIN |
| PLOD2 | GSN |

| | |
|---|---|
| ISCA1 | FLJ16331 |
| UCK1 | IQWD1 |
| SPNS1 | ADRA1B |
| SP9 | KCNK17 |
| RUNDC1 | HOXA10 |
| GPR160 | RDBP |
| 5031414D18I | ACRBP |
| SLC14A2 | SAMD1 |
| D10JHU81E | TA |
| CD86 | KLHDC6 |
| SRSF3 | LOC284013 |
| LSMEM2 | LOC283932 |
| LFNG | SILV |
| ICAM2 | ATP6AP1 |
| AUH | SCGN |
| SLC25A39 | CPT1B |
| MBD2 | FLJ20200 |
| TNFAIP3 | CYBASC3 |
| PSRC1 | MYH10 |
| ALAS1 | RPS26 |
| FOXO1 | SLC36A4 |
| MRPL37 | TNK2 |
| NDUFV2 | KIF5A |
| PDK4 | PKP4 |
| DUSP10 | SEZ6L2 |
| FARS2 | MGC16028 |
| PKP2 | CENTD3 |
| CCDC28A | SBBI54 |
| AMPD3 | C10orf45 |
| PIAS1 | C2orf24 |
| LRRC39 | IQCG |
| GLTPD2 | SIMP |
| GYLTL1B | SEMA7A |
| TMEM134 | MADD |
| STAT1 | TUB |
| GCLC | KLF12 |
| ZC3H14 | LOC128439 |
| CAR9 | PCDHAC2 |
| COL8A1 | KIAA0406 |
| OLFR284 | MCAM |
| COLEC12 | MGC51082 |
| RPP30 | GABRA2 |
| BLVRB | GALNT7 |
| REEP1 | OBSCN |
| GTPBP2 | MYL1 |
| UBE2T | MGC32871 |
| RPH3A | PCGF2 |
| CWC25 | PCANAP6 |
| PRELID2 | FLJ13391 |
| ELK4 | IQCD |
| MKKS | MDM1 |
| PROCA1 | FCAMR |
| BANF1 | CD9 |
| UBE2QL1 | PPP1R10 |
| FAM13A | PTX3 |
| SLC22A3 | NXF1 |
| BSG | TP53BP1 |
| GADD45B | KIAA1285 |

| | |
|---|---|
| TRPV1 | DC12 |
| FAM84B | CAMK2N1 |
| BBX | C11orf8 |
| UCP2 | NUMBL |
| DTX4 | MST1 |
| BNC1 | H2AFX |
| FOXG1 | FBN1 |
| FGFBP1 | INHBB |
| P2RY14 | BAT3 |
| ALS2 | MCEE |
| PABPN1 | HOXD1 |
| NDUFB3 | AIF1 |
| IGSF21 | ALDOA |
| SLC22A5 | HIST1H2AI |
| SLC25A42 | FLJ44635 |
| FKBP8 | LASP1 |
| FMNL1 | KIAA1853 |
| PTCD2 | SKIV2L |
| WFDC6B | TBX15 |
| MMP11 | KCTD9 |
| B3GALT2 | LIN28 |
| MIPEP | FAM11A |
| CDK18 | MGC29649 |
| UCHL3 | HEBP2 |
| PPARD | GABPA |
| FSHB | HCNGP |
| ANXA2 | FLJ20457 |
| MCF2L | USP37 |
| SPEG | LOC255374 |
| TBC1D24 | PSME1 |
| STK10 | STAG3 |
| GREM2 | MVP |
| ARHGAP20 | EIF4G1 |
| ABCD3 | C1QL4 |
| HDDC2 | CREB5 |
| VPS26B | LOC90355 |
| INPP5F | C14orf111 |
| MYBL1 | AP1GBP1 |
| PAQR7 | ACE |
| INPPL1 | SLC35A4 |
| PDZK1IP1 | MLX |
| ACOT6 | PCGF4 |
| CHGB | PTE1 |
| TMCO6 | FLJ36268 |
| FAM73B | SLC25A25 |
| SP7 | LOC92154 |
| CES1E | TAX1BP1 |
| CDC27 | RARSL |
| MDGA1 | MGC26816 |
| COPS6 | MGC30208 |
| 2310030G06 | PRKAA2 |
| CNN2 | GIT1 |
| BRI3BP | C14orf35 |
| GPC4 | HNRPUL1 |
| WNT1 | LOC55974 |
| DBP | FAM35A |
| LMO7 | DPF3 |
| SOGA1 | MOV10 |

| | |
|---|---|
| COL4A2 | LOC348094 |
| NGP | LOC200383 |
| BPGM | RARA |
| CDRT4 | FUT1 |
| SLC12A7 | NPHP4 |
| TCF7L1 | DLG2 |
| HMOX1 | PPP1R12B |
| CITED2 | HH114 |
| DTX2 | ANAPC11 |
| GRM3 | SERTAD3 |
| CD302 | WDR46 |
| STARD8 | RORA |
| FAM126A | LSP1 |
| PPP2R4 | DEPDC2 |
| SIAH2 | SLC22A17 |
| PFAS | CCR1 |
| CCDC103 | C21orf127 |
| CSDC2 | HNRPDL |
| ABHD5 | DUSP6 |
| MKNK2 | NUP188 |
| TMBIM1 | RAB3 |
| VSTM2B | CNIH2 |
| EFNA5 | CPEB2 |
| FCGRT | OTX2 |
| NXN | SMARCD2 |
| IGF2BP2 | ELAVL4 |
| URGCP | DERPC |
| EMB | SYNC1 |
| PNPLA3 | DKFZp434O0527 |
| MAF1 | BCORL1 |
| CHRNA6 | TPP1 |
| KRT80 | RREB1 |
| ACADM | GPR115 |
| GIPC2 | VEGF |
| SLC22A23 | SEC24C |
| DUSP3 | IRF2BP1 |
| ANKMY1 | CIC |
| IRX6 | TNPO3 |
| ANGPTL4 | PPGB |
| HHEX | FGF9 |
| MPZ | SLC1A4 |
| NAMPT | MGC33951 |
| HECTD1 | HINT2 |
| SLC31A1 | DOK3 |
| HACL1 | DNCLI1 |
| RAD23A | EDA |
| ACY1 | JUNB |
| SFXN1 | LOC56901 |
| RNASEH2C | 76P |
| FAM102A | PMP22CD |
| TECTB | TP53INP1 |
| IRAK2 | FLJ45300 |
| GHITM | CAP350 |
| NDUFB7 | ZBTB9 |
| FOXN3 | LOC196996 |
| TXLNG | HRMT1L4 |
| NTRK2 | NTN1 |
| MFGE8 | WNT10A |

| | |
|---|---|
| SP110 | NEUGRIN |
| YAF2 | HSPBP1 |
| SH3BP2 | A1BG |
| D130043K22 | CSAD |
| PXMP4 | LOC541565 |
| TFB1M | PARCC1 |
| HDHD1A | ABCF3 |
| DCAF4 | CCL28 |
| ATP9A | ATPAF1 |
| LCLAT1 | C1orf84 |
| TSKU | CEI |
| PDLIM1 | POLR1D |
| ETV3 | SNAG1 |
| UQCR10 | USP26 |
| ISLR | STEAP2 |
| CLIC4 | HNF4A |
| TLE3 | hSyn |
| DDX31 | RNASE11 |
| TULP1 | TSEN34 |
| 1700017B05I | KHSRP |
| PPP1R14B | DDC |
| ALOX5AP | SEMA4G |
| MESDC1 | MGC15429 |
| ADHFE1 | DST |
| GNAZ | RAB5C |
| DDI1 | RIT1 |
| ZFP777 | POLR2H |
| GYS2 | SCML1 |
| MMP14 | WTAP |
| SERPINI1 | MLF2 |
| PLAT | LOC389816 |
| NUPL2 | DSG3 |
| HDHD3 | SPBC24 |
| POU4F3 | AKT1S1 |
| TDRD3 | C1orf117 |
| PPP2R5B | C14orf4 |
| CALHM1 | LCN7 |
| KLHDC3 | RAD17 |
| GNAI1 | RILP |
| SLC25A4 | LGI3 |
| CYB5B | C6orf136 |
| BTN2A2 | WFIKKN2 |
| NDRG1 | BSCL2 |
| SESN1 | ANKRD12 |
| SDHAF2 | SDHC |
| CDHR4 | SNIP |
| TMED8 | FLJ13236 |
| ARHGAP1 | LMO3 |
| GBE1 | BAI2 |
| PIP4K2A | BACH1 |
| RASGEF1B | FLJ25660 |
| SOD1 | KIAA1924 |
| LDHAL6B | MGC50811 |
| AFAP1 | EWSR1 |
| PACS1 | C20orf70 |
| FSCN1 | MRPL21 |
| TMEM136 | USHBP1 |
| SIRT5 | PPP1R16B |

| | |
|---|---|
| MSRB2 | ARMET |
| ZFP36L1 | C1orf8 |
| SRPX2 | C10orf94 |
| ACSBG1 | RASAL2 |
| CBR3 | ZNFN1A2 |
| POLR2F | PLCD3 |
| WDR74 | TMEM30A |
| ST6GALNAC4 | NEDD9 |
| MCM7 | ELF5 |
| ATP5G3 | HOXB9 |
| METTL21B | GCAT |
| LYVE1 | FBXL11 |
| APOH | RBM8A |
| PDX1 | NOB1P |
| ABCB9 | FLOT1 |
| TSC22D2 | LOC90580 |
| ASPG | SLC25A28 |
| STAB1 | ZNF462 |
| ASPDH | MGC33600 |
| CFL2 | FLJ25328 |
| RASA2 | SLC6A9 |
| SLC22A15 | FOXP1 |
| STX18 | MGC21688 |
| KCNA3 | POFUT1 |
| DBI | PLEKHA4 |
| BTBD1 | CXXC4 |
| LPIN2 | LOC220594 |
| TMIGD1 | WDR18 |
| ZHX2 | SCRIB |
| PDP2 | WDSOF1 |
| CD28 | PITX3 |
| SMARCA4 | HPX |
| RPS6KA4 | LMX1B |
| ARMC9 | PFN1 |
| C1QTNF6 | MGC20983 |
| TTC8 | JUB |
| CLSTN3 | FLJ45121 |
| NFE2L3 | FLJ33387 |
| CEP55 | IMPA3 |
| PALMD | STAC3 |
| HS6ST1 | C11orf31 |
| TTC39B | KCTD12 |
| ENPEP | CBX5 |
| SH3D21 | REM1 |
| CTSE | HS322B1A |
| TIMM23 | C1orf94 |
| OAS2 | ZNF436 |
| GABRD | FLJ11795 |
| EPYC | SCAP1 |
| BIN2 | DGAT2L3 |
| DEGS1 | PEX5L |
| CD74 | PLA2G4B |
| LIPG | ING1 |
| FAIM | RASGRF2 |
| KRT16 | TRPS1 |
| 1110032A03I | CRYGA |
| KLF17 | SLC14A1 |
| UBXN11 | SET8 |

MGST2
SECTM1B
SLC25A11
FAM154A
STARD4
OSBPL1A
PLCZ1
GPR35
GYK
PLEKHH1
RPL3
KLB
ACSL1
TTI1
PIK3R2
TNFRSF21
CDC42EP5
SLC6A8
CDC45
TMEM38B
MC2R
SMPD3
TAZ
NFE2L1
OLFR71
STIP1
FOXS1
NT5DC2
DIO2
LRP2
CYTIP
CHST7
CCT3
SLC25A20
IER5
HSPE1
MAPK13
EIF4EBP1
APLN
C2CD2L
SNRK
GUSB
PMPCB
DMRT2
NID2
HYLS1
GLDC
CD1D1
NUDT16L1
CHEK2
ZFP365
DRD4
PRKRIP1
ARL5C
ADH4
KRT18
PI4K2B
NTSR2
LHX2
CXCL16
GAB1
FLJ22635
NXPH1
KCNN4
LOC158160
FLJ25801
DLEU1
CLCF1
GIT2
TNFAIP8L2
NUDCD1
CHCHD4
MGC26733
FTSJ2
DPP10
C21orf58
SCNM1
ARHGAP22
NPAS3
FLJ20171
PYDC1
C3orf21
PGEA1
FANCB
DKFZP434H132
TOB1
EEF1G
FLJ22104
DNAJC13
FLJ13154
U2AF1L3
PFDN2
HIST2H3C
PRR3
ProSAPiP1
IL11RA
SNRPD2

MCM4
TCEA1
RNF111
CHST12
CLPX
MRPS36
RAB34
PJA1
CHMP1A
GM8140
APOC2
PHLDA2
BZW2
PTPN6
TST
LYSMD3
ERLIN1
GPX8
CHCHD10
GLRX
INHBE
CCDC85C
RHOG
CYB5R3
HEY1
ITGAX
CDK2
GPAM
CDC37
PLIN5
PRPH
FBXL7
LRRC8B
SEC14L2
IL1RN
MBNL3
TRP53BP1
YIF1A
ADORA2B
MAFB
TMEM218
PHEX
GRINA
TFPI
3110043O21RIK
H2AFV
TMEM87A
TBC1D22A
HIP1R
CYP17A1
TOMM70A
SCG3
DGKA
PQLC1
KRT19
ENO1
SBDS
GABRB3

CXCL10
GIPC3
DET1
AZIN1
SLC7A7
RNPS1
MB21D2
NEURL4
ISOC1
SLC25A10
TMEM64
ARG2
MKI67
ADAP2
FFAR2
PEX5
LENG1
ZFAND5
HAO2
FH1
MAFG
MAP1LC3A
NUDT13
CCL6
TOMM5
DAK
ZCCHC2
PAXBP1
PLA2G4D
LOX
FIZ1
YARS
ELOVL1
UBE2E2
LAMB2
DTD2
PBXIP1
OLFR285
MARCKSL1
IMMP2L
RAB11FIP5
BAG4
CDKN1A
CHCHD6
KLHL34
A430005L14RIK
CAT
PRL2C5
CUL7
DIO1
CPOX
SPSB1
RNF10
RARRES1
TPM2
PPP1R11
SIM2
TMEM120A

FAM170A
SLC2A2
LRG1
WDR48
PIK3R1
NRXN3
ACACB
MAF
HOXC11
TRAK1
ADCY4
CSTB
ADM
HILPDA
PREPL
RHPN2
DOC2B
STAMBPL1
PTGR2
CRYL1
ACP6
PPP1R1A
SELM
ETFB
TMIE
HADHA
IDH1
SEMA4A
GM11397
PLA2G16
RBM39
HDAC10
ZFYVE27
ELMOD3
FXYD1
LMOD2
SHISA7
VSIG2
RGS17
RUFY3
CLDN1
PCTP
ABCC1
MRPL51
TMEM135
SLC27A2
BCL6B
HAX1
ZFP467
GDF1
REN
NEK9
KDM2B
G6PC
ACTA1
OVOL1
DDO
NHLH1

SLC25A45
CTSA
TOMM40
UBFD1
TCF23
TRIM46
NOBOX
MMADHC
PIP5K1B
TEKT5
MBOAT7
LCAT
HOXC8
AIG1
ARHGEF25
ST6GAL1
APBA3
SLC6A20A
FBXW17
APEX2
FAM181A
SLC24A3
CYP4A31
HMGCS1
DEDD2
HAUS7
ZBTB3
MRPL23
ORC4
ITGA7
CNOT3
MTIF2
HIBADH
TSSK2
MPRIP
TSPAN4
TXN2
ZFAND2A
ALB
LY6D
CPT1A
C1QA
LMO1
CEACAM1
SREBF1
GRIA4
TMEM132D
EFR3A
SUCLG2
LOC102639054
EPAS1
SCNN1G
COL8A2
AMPD2
IFNAR2
CYP8B1
OASL1
LPIN1

SLIT2
PDE1B
TNP1
CCP110
ITGAE
ODF3B
GAPDH
SUCNR1
ELK3
WFDC1
COMMD6
FCGR1
SLC39A1
NARS2
LIN28A
DHX58
LIMK1
ZFP191
FAM20C
TSPAN17
GCN1L1
CLEC10A
AMOT
HOXB7
TMTC1
TIMD4
GSTM2
CTXN3
ADAM11
0610010K14RIK
PCOLCE2
ZFP40
ATP6V0E2
PEAR1
KRT79
FABP2
ME1
CDA
PGS1
ZP2
KDSR
MCM8
SETD4
TEX11
SLC4A4
ACOT8
BHLHE41
GAREML
SMAP2
FAM101B
PZP
KEG1
GJB4
PEBP1
MRPS18B
RPL14
PCSK7
FBXO2

AOX1
RMND1
AMIGO2
TMEM26
LRTM2
LTBP3
ZFP687
CTTN
EVPL
ABTB1
KCNJ8
PASK
SLC16A1
EHHADH
RANBP9
USP8
TMEM147
LCN8
PIPOX
LNX1
SOX18
1810026J23RIK
INSIG1
SYNGR1
CFD
NR1D1
APOC1
PRAMEF8
TMEM37
ITGAL
2010111I01RIK
ERC1
WDR73
YPEL3
THEMIS
MLYCD
SCD2
NDUFA5
SCO1
ZBTB40
SDHA
NCLN
ALPL
TMEM206
CACNA1G
PDLIM5
SAT1
ZFP703
BRWD1
THNSL2
RCBTB2
C1QTNF1
AS3MT
CYP4A10
LETMD1
UQCRC2
HYAL1
PVR

OLFR1328
ISCA2
ARTN
EYA2
BEAN1
BCL2L14
S1PR3
MAGEA4
GNB3
ANKZF1
TMTC2
L3MBTL1
LBR
CBFA2T3
PGK1
WLS
AGPAT9
PCP2
KLF3
CDC25A
ANKRD37
NAT14
RCN3
RRN3
B3GNT5
PTGER3
COBLL1
CHPT1
TEX264
ERGIC1
RPTOR
BARHL2
E2F8
ADH1
XPNPEP2
EIF4EBP2
TLX3
PVRL1
HSPB6
IER2
WASL
MACF1
D930015E06RIK
CHRNE
TRAF3IP2
MDFIC
MED20
EHF
ADD3
VMN1R158
AMICA1
RAP1A
ERI1
TRIO
TC2N
6430573F11RIK
ROGDI
MRPS35

C1QC
RASSF3
CILP
ATXN7L2
ACOT1
RDH13
HOXC13
SLC15A5
SLC25A37
HMGB3
NLRC4
CTNNBL1
ECHDC1
MID1IP1
NDUFA6
CPSF4L
GM867
DNASE2A
NDUFAF3
KCNE1
PHLDA3
DSP
RAD51B
GALK1
NUP214
SUPT16
IRF1
FBXL6
DPYSL2
SOX13
SSH2
SDHB
SOX4
PRAM1
NGF
TSPAN3
AK3
DNAJB1
RHOF
USP20
GPR101
NRP1
ALOX5
PLIN4
TUSC5
MDGA2
FBP2
JAM3
FASN
MRPS14
LAPTM4A
PHLPP2
BTNL9
PRDX6
GPNMB
TTF2
FGFBP3
NXPH3

SLC25A26
DOLK
ACVRL1
GPX7
TGM2
PSMA1
FCGR2B
ACYP2
2700049A03RIK
NFATC4
DEFB1
TMEM60
CRYM
YBX2
ARPP21
NR1H3
CIB2
ABCG1
RGS3
SQSTM1
AADAC
CCR9
DSCAML1
SNRNP70
SLC35F6
PRICKLE3
UAP1L1
ACAT1
SERPINB1A
NEURL2
MEOX1
PLK3
RALY
PHKA2
ATF3
UCHL1
FBXO22
MRVI1
DGKH
GJB6
TAGLN2
SLC16A7
NDUFA11
PUSL1
CES2F
RASL10B
MRPL19
TAPBPL
PRKAR2A
KLHL23
SYT10
CNOT4
PEX14
KLF16
PGRMC2
PLA2G2A
PLTP
CHST1

LGALS1
THOC2
XIRP1
DNAJA4
MINK1
SLC25A1
EZH2
RAP1GAP2
DOCK8
EXOC6B
PLEKHB1
GRB7
SRI
SOCS6
SLC25A5
YWHAE
WDR1
AFG3L2
ZFP575
KRTAP7-1
NEUROG3
ADAT2
CNKSR3
MGST3
HBEGF
CDKN1B
R3HDM4
NDUFS8
RASAL1
ZC3H3
DDAH1
ABCA1
MGAT1
TNFSF10
TIPARP
SLC8A3
DNASE1L3
CRYAB
MRPL47
MUC1
RASD2
TMEM143
CAR4
SLC40A1
RNF34
ACOX1
SNHG11
CPLX1
CENPH
LRRC17
ANAPC13
FAM183B
SLCO3A1
HSD17B11
IDH2
HSD17B4
SEC31B
ZNHIT1

SLC12A6
EFHD1
4930402H24RIK
COPZ2
ELF3
CPSF3L
NUDT7
NOP2
NNAT
ANGPTL7
ADAMTS9
GPCPD1
BCL9
MRPL28
STIM1
PIGV
CDK2AP2
CXCR2
LETM1
APBB1
CAMK2D
ALG14
IFITM5
DNAIC1
LIPE
PODXL2
CELF6
SLC25A44
ADCY5
4921511H03RIK
ESYT3
8430408G22RIK
SFSWAP
FGD4
CLP1
ZFP524
PTGES3
KIF16B
CASP8
CS
BNIP2
NRP2
CPSF7
MED10
CDH2
PTRH1
2810004N23RIK
DEGS2
COX17
ERMP1
GTF3A
MRPL54
POU4F1
SIPA1L2
CCNG2
RCAN2
EMG1
NCEH1

WDR5B
HIRA
MERTK
APLNR
CITED1
SLC47A1
ORMDL3
CORO2B
PAN2
TECR
CCRN4L
AKNAD1
RNF144A
OLFR1259
CBR4
SLC1A5
CCL7
LPXN
SPEM1
6820408C15RIK
TNNC1
GFI1B
PUS1
PEX1
OLFM1
TUSC2
2610034B18RIK
FADS2
ENAH
HSPA4L
SMAGP
TRIM55
STBD1
TAC4
FOXO6
CLDN7
PARP14
PTGFR
TRIM37
ABHD15
AFP
CRY1
OLFR90
MSX2
NUP62-IL4I1
ABCC4
CUZD1
FAR2
PDRG1
FAM69B
ZCCHC24
AU018091
CTF1
PIM1
S100G
NR5A1
THBD
DNAJC7

FBXO40
SERPINA3N
CHKB
PARP11
NOL3
CD180
ACTR6
MYBPH
MRPS18A
LYRM5
CHCHD7
PPP1CC
SERINC2
TM7SF2
CDCA7L
JMJD7
HDGFRP3
LEPR
CCDC109B
MTOR
SRP68
RAB9
SV2B
NDUFS2
SLC19A3
ZFP704
WT1
SLC2A8
ILK
SLC25A22
STK38L
SLC22A1
MRPS7
AOC3
MTHFD2
FABP3
NPTX2
TRPV2
SLC27A1
RALGAPA2
CAMKK1
RORB
OAS3
ANXA6
LRIG3
P2RY1
VAPB
42432
AP2A1
C2CD4C
FZD7
AFF4
TIFAB
NCCRP1
MRPS33
STOM
AIM1
RAMP1

SMARCA5
CLDN23
GAS2L3
UNC79
H2-D1
CES1D
MYH3
PDE8A
RSPH1
PLA2G4A
SMOX
ARPP19
FKBP10
PRUNE
SMAD7
VEGFC
ITGB6
DBT
SDC4
SLC1A2
ZBTB16
CALR
HIGD1A
WARS
LTB
IL4RA
RGL3
FAM149B
NAA50
ASPA
ABCC3
CERS1
AKR1B8
MAGIX
PIP5K1A
ITPKA
CARS2
BTBD2
BEST1
MMACHC
REV3L
TWF2
RHBDL3
ZFP628
CYC1
EDNRB
CD27
DOXL2
RHEBL1
PEX13
DISP1
SGTB
3110002H16RIK
RCVRN
CHUK
RAB33A
ZDHHC12
DENND4B

PCX
PDK2
SRGAP2
SLC38A2
SYNJ2
STK32A
TRIM26
ST3GAL6
KRT36
SPIN4
TRPT1
NUDT14
CAP1
DUSP1
SLC44A1
GFM1
PCK1
NPR3
KBTBD11
SULT2A1
PRPS1
NDUFB5
LIPA
CIDEA
CDC42EP2
SETD6
GAMT
OXR1
SOX17
DPYS
MRC2
GARNL3
PNPLA1
SLC6A13
MIDN
FRAT2
AKR1C19
VTN
BAD
NAA10
LRPPRC
NDUFAF7
CIDEC
EBF1
CYP4A12A
MRPL35
CYP2E1
ACO1
RBX1
1700034O15RIK
GRHPR
MEGF9
MYO18A
HTRA3
GABARAPL1
ARRB1
TINAG
PFKFB3

RUNDC3B
SCAMP5
ECSIT
LRRC15
IL17RB
TFF2
RNFT2
GSTT2
RNF168
TRIM69
MTERF4
CYP2F2
CRYBG3
USP2
CISD2
ABCG4
CALY
COQ3
PLCD4
RRBP1
UBXN2B
PRDX5
VDAC1
VDAC3
CASP7
2200002D01RIK
KCNJ10
RDH16
TOM1
LACE1
PRMT6
D630045J12RIK
CDC20
KCNE1L
TMEM106A
ACSL5
NDUFB6
ADRB2
TRP53INP1
TYSND1
NMT2
CTSD
RETNLB
RRP1B
SLC4A7
SGMS1
WDR64
SEMA3B
TUBB3
THOP1
SCARB1
2300005B03RIK
ADSSL1
URAH
S100A1
MRPS26
SLC7A10
RBPMS2

AK2
ITPR2
THRSP
SPRR1A
LRRC10B
SH3TC1
BHLHB9
GRB14
NTRK3
OMA1
RFC4
BCL2L13
FMO1
CCDC30
NUP50
TMEM63B
SEMA4B
KAT2A
CSNK1A1
2310007L24RIK
MMEL1
MFSD1
1700112E06RIK
PUS7L
PEX6
UBAP1
DHRS3
DYNLRB1
PBK
MR1
BC048679
EXOSC5
CHODL
TNFRSF12A
CCDC62
ATF4
IFRD1
IL12A
GDPD1
ARNTL
9130401M01RIK
MTNR1A
SIGMAR1
TSPAN5
PUS10
CCL2
AMIGO1
ATAD3A
ACTN2
KRT81
POPDC2
NIM1K
TRIP13
MAZ
DFNA5
FAM26F
ASGR1
ACADS

PVRL2
IL6
SNAI3
MED30
ANXA5
EPHX2
CORO6
KIF3B
ANGPT1
CPT2
GLMP
CHAC1
LSS
TOR1B
EPOR
ZMYM1
EPB4.1L4B
ARHGEF40
5031439G07RIK
UBE2D1
IDH3A
HSP90AA1
ASB15
ANGPT2
SEL1L
YWHAB
ACSS2
LY6G5C
FGFRL1
PLIN1
SUCLG1
EIF4B
SSPO
SEC23A
MAGEH1
PRKCE
ITGAM
RILPL2
PGM2
ROCK1
TUBE1
RGS2
TAGLN
CREB3L2
RTN4
GFM2
NDUFB10
CCNO
RXRG
IVNS1ABP
CD163
PGP
RGS5
NDUFV3
OPLAH
ACSS1
ST3GAL2
ABCG2

MRPL18
AKR1B3
FBXO21
TUBA4A
ACRV1
HPD
CLSTN1
LANCL3
MYLIP
RPIA
PRDX3
HNF1A
LMAN2
CHCHD3
MFAP4
SLC25A47
CES2G
SLC11A2
H2AFJ
TMEM45B
DHX32
DENND5A
TMEM150C
STXBP2
SEMA5B
BDH1
COQ6
COTL1
MS4A1
DOCK9
SCYL2
MYH14
GADD45G
UROD
DNMT3L
DDX28
FITM1
RBCK1
ZDHHC8
PLXNA2
MGLL
FBXL15
CYP4X1
PLCG2
BCMO1
CRYGD
DDIT4
NUS1
ITGA2B
CLDN5
NCS1
UGT1A9
ATP5L
ATP1A2
CYP1A2
ONECUT1
NUDT9
BACE1

HERPUD1
CABP4
BCL2L11
RBP4
ATP5O
METTL9
TIMM9
CDO1
UGDH
MAP2K5
LGMN
SLC5A6
SDPR
SEPP1
SRXN1
LAMB1
IGF2
SERPINA7
NUCB2
RARRES2
MEIG1
STARD10
HOMER2
ANAPC2
KCNS1
NOD1
1100001G20RIK
ADAMTS1
JMJD6
CTSL1
PRKG1
INA
DGKI
FAM136A
NFIX
SLC16A6
GRID2IP
AKAP1
HIBCH
SUV39H1
SLC7A5
AIFM2
TIMM10
AK4
NLN
SORBS1
ETFDH
LY75
PCSK4
GM1673
UMODL1
CYGB
NME7
NQO1
HSPA9
AXIN2
DMBT1
KCNAB2

SPNS2
1700001C19RIK
GSTK1
MAP3K2
SLC9A6
TSPO
HOXC9
UCP1
RAB3D
FOXA1
TBPL1
ARRDC3
VNN1
OLFR456
SLC16A13
SDC1
PLCL1
CACNA1A
TEX22
ALDH3A1
MUL1
GGT7
GALNTL6
ACSM1
PSAT1
IL1F6
OSR1
PTGR1
FOS
CNTRL
MRPS5
AGPAT2
CREG1
HSPA5
RETN
TNFSF13B
SLC6A19
CMPK1
OGFOD3
CYP27A1
PAFAH2
SIGLECF
NPEPL1
H6PD
GNPNAT1
SAP30
TMEM190
IGDCC4
SECISBP2L
HTATIP2
SPDEF
PEX3
PAMR1
FXYD7
PABPC4
HADH
TRPM1
CCKAR

SASH3
FAM109B
ABCB10
IDI1
KCTD6
DDN
SPACA4
GLUL
STKLD1
TRAF4
HK2
ELOVL6
WNT8A
CYP4F15
EPHX1
DLX3
ARAP2
DNAJC2
GADD45GIP1
SLC27A4
PPIF
EHD3
COL14A1
PRR15
TNFAIP8
EFEMP2
ABAT
TRIM54
CAV1
DGAT2
TMEM35
HECTD3
STRBP
ABHD3
GTF3C4
LGALS9
SCARB2
IQSEC1
CYP4V3
BAG2
PNMAL1
PLBD1
ANKRD2
RAD54L
HMGCL
ALX4
TMEM164
SLC9A1
THBS2
ASNS
PPM1L
ISG20
TTYH2
GJD2
GATA5
1600014C10RIK
TEC
HSP90B1

KCNK15
SMOC1
ABCB4
COX8B
TCEAL5
GPR146
SORD
MFSD4
SPC25
SLC25A33
FAH
H2-AB1
RCE1
UBE4B
TIMM8B
TRIM44
FRK
USH1G
STAP2
PPAP2B
DNTTIP1
CRNN
RASD1
CABP2
TRAF1
GRN
FAM98C
SLC2A4
HOXA4
TRIM32
LDLRAD3
LRRC59
C1QB
BMYC
ADCY6
GLYAT
TGFA
NPR2
TMEM170B
PRLR
MATN1
AP3S1
3110057O12RIK
RTP4
INMT
LAS1L
FLAD1
NPPC
ERP44
PKN1
ITM2A
SPON2
RHBDF2
CEBPA
ACSF3
JMJD1C
ALOX12B
TSC22D4

COX6B2
OLFML2B
S100A10
THOC6
WDR76
MRPS2
EFTUD1
ACBD4
CDKN1C
SIRT3
AQP1
ST6GALNAC6
ACER2
THUMPD2
ACTB
APCDD1
TRMT12
CMBL
CNKSR1
RAB30
PCDH9
OSBPL11
GBF1
ACVR2B
IYD
ZSWIM7
SERPINE1
DNASE1L2
ZMYND12
UBQLN1
EXTL1
POU2F3
GBP3
ATAT1
ARC
PPP2R2D
TIMM44
LRRC51
TMEM128
ILVBL
RNF186
OXNAD1
SMIM20
2210016F16RIK
TMOD1
PLEKHH3
PRELID1
RSAD2
FAM162A
INSR
INTS4
PLLP
GLRB
GALE
AMN
CIZ1
NT5C2
MSLN

LMTK3
WWTR1
CIB1
SLITRK1
SLC45A3
OXSR1
TNFAIP2
RAD51D
MTSS1
MEST
ADPRM
DLD
RGP1
MMD
ADAD2
KLF15
PCDHA1
CUL2
DENND4C
PKM
MORC4
ETV5
LY6E
SCUBE2
S1PR5
GSTO1
COX6A2
NRCAM
FABP4
LSM1
IDUA
CSRP2
LYRM4
HIPK2
CCND2
ADIPOR2
PEX11A
CDC34
PKDCC
RNMTL1
SYT17
ECHDC2
RTKN
UPP1
CLEC4G
APOA1BP
CISD1
TMX2
GPRIN3
PPM1F
CRTAC1
PCSK5
GMNN
NDUFA9
SUDS3
RNF149
CDCP1
EGLN3

PEX19
PNLIPRP1
ERCC6L
GPX3
UNC119
EDN2
NOV
PSMD7
SHISA2
CCDC121
SSX2IP
ST8SIA2
LRP4
TMTC4
SNRPB
TAPT1
ENC1
IRF8
MICALL2
SOX11
POP5
CGN
TCEA3
TLCD1
LPCAT3
SF3B4
VOPP1
GTF2IRD1
LTC4S
SMARCA2
ACOT4
STC2
COL6A1
FOXD1
PRSS8
WNT4
GPRC5B
GNG13
F11R
HSD11B1
MMRN1
VIPR1
ACTA2
DERL1
SP5
TIMM8A1
4930538K18RIK
LAMA4
F3
PPFIA4
MAP4K4
CYP2D22
PLEKHM2
PAQR9
DLAT
KIF21B
SYNPO
KCNJ5

LMNA
TRIT1
HOXC6
CASQ2
PRICKLE1
PEG10
CCDC92
KLF10
FOXO3
TSNAXIP1
FTCD
PARD3B
PHOSPHO1
DLX1
LPAR1
RAG2
2010001E11RIK
PPA1
XIRP2
POPDC3
GEM
DOK7
TFPI2
NANOS1
PDHB
SHKBP1
CSF1R
JUN
NAGK
MYL2
KDM3A
ATP2A2
RBP1
MYOM3
RETSAT
ADAM15
FAM110B
CALHM2
AOAH
FBXO32
DCAF12L1
PNN
TGFBI
CLPP
PSMB10
PAM16
2610002M06RIK
PPP1R3G
TRIM25
SDCBP2
NUP62
GATM
RASA3
HNRNPL
PGD
RUNDC3A
ITPRIP
SPINT2

E2F3
ACOT3
RNF128
ABHD6
CUTA
WAC
SLC25A48
CD274
LRRD1
ATPAF2
PTPN13
ECD
FDPS
SLC52A2
DAPK1
SUCLA2
VAV3
MTMR11
IDH3B
SUOX
SELENBP1
NDUFS7
PRKCA
NPC1
ALDH9A1
CD40LG
STK17B
B3GAT3
ATP8B1
3010026O09RIK
MAT2A
LRP11
TRPC3
PGPEP1
MBOAT2
PRODH2
SEMA3C
HCAR1
DPYSL3
SMYD1
2310061I04RIK
S100A13
PAK2
MNX1
CSF2RB
PARM1
IFI44
SMAD5
GARS
FAM49B
PNPLA2
BCKDHB
RETNLA
KRT23
SERHL
PNKD
HDAC6
UBC

OTOP1
PSMB9
SNCB
ADRBK2
TNNT1
GPR15
GHRH
CBR1
GSTM7
RXFP2
ZNHIT6
LIPT2
PIK3IP1
ADIPOQ
RXRB
